# Rapid changes in plasma corticosterone and medial amygdala transcriptome profiles during social status change reveal molecular pathways associated with a major life history transition in mouse dominance hierarchies

**DOI:** 10.1101/2024.05.30.596669

**Authors:** Tyler M Milewski, Won Lee, Rebecca L. Young, Hans A. Hofmann, James P Curley

## Abstract

Social hierarchies are a common form of social organization across species. Although hierarchies are largely stable across time, animals may socially ascend or descend within hierarchies depending on environmental and social challenges. Here, we develop a novel paradigm to study social ascent and descent within male CD-1 mouse social hierarchies. We show that mice of all social ranks rapidly establish new stable social hierarchies when placed in novel social groups with animals of equivalent social status. Seventy minutes following social hierarchy formation, previously socially dominant animals exhibit higher increases in plasma corticosterone and vastly greater transcriptional changes in the medial amygdala (MeA), which is central to the regulation of social behavior, compared to previously subordinate animals. Specifically, social descent is associated with reductions in MeA expression of myelination and oligodendrocyte differentiation genes. Maintaining high social status is associated with high expression of genes related to cholinergic signaling in the MeA. Conversely, social ascent is related to relatively few unique rapid changes in the MeA. We also identify novel genes associated with social transition that show common changes in expression when animals undergo either social descent or social ascent compared to maintaining their status. Two genes, Myosin binding protein C1 (*Mybpc1*) and μ-Crystallin (*Crym*), associated with vasoactive intestinal polypeptide (VIP) and thyroid hormone pathways respectively, are highly upregulated in socially transitioning individuals. Further, increases in genes associated with synaptic plasticity, excitatory glutamatergic signaling and learning and memory pathways were observed in transitioning animals suggesting that adaptations in these processes may support rapid social status changes.

**Author Summary:** We investigate how male CD-1 mice behaviorally, physiologically, and molecularly respond to changes in their social hierarchies. We found that mice rapidly reform social hierarchies when placed into novel social groups. Formerly dominant mice experience the greatest increase in stress hormone levels and the largest shifts in gene expression in the medial amygdala, a brain region crucial for regulating social behavior. Specifically, we observed reduced expression of genes related to myelin production and maintenance as well as cholinergic functioning in formerly dominant males who socially descended. In contrast, formerly subordinate males who socially ascended displayed fewer genomic changes. Moreover, we identified a set of genes that change expression when animals transition their social status regardless of whether they rise or fall in rank. These social transition genes provide new insights into the biological mechanisms underpinning social plasticity. Our findings demonstrate the distinct molecular mechanisms that are shared between socially ascending versus descending mice, as well as those that are unique to each type of social transition.

## Introduction

Dominance hierarchies are a common form of social organization across diverse taxa (Strauss & Shizuka, 2022). Individuals establish social ranks based on the outcomes of agonistic interactions, with more dominant individuals monopolizing prized resources such as food, mates, or shelter. Stable social hierarchies emerge when costs for continued conflict are high, meaning that all individuals benefit from establishing resolved social dominance relationships (Chase, 1982). However, individuals must be able to dynamically alter dominant and subordinate behavior in response to social and environmental challenges and opportunities (O’Connell & Hofmann, 2011). For example, animals may seek to socially ascend a hierarchy when dominant animals are successfully deposed by lower-ranking individuals or removed by predators. Conversely, animals socially descend when animals of lower rank become more aggressive or more competitive animals join the social group. Such transitions in social rank are associated with significant modifications to individual behavior and physiology (Dwortz et al., 2022; Friesen et al., 2022; Maruska et al., 2022; Milewski et al., 2022; Snyder-Mackler et al., 2016). Although individual ranks may change in hierarchies, periods of instability tend to be relatively transient with hierarchical structures rapidly reestablishing to maintain structural stability (Chase et al., 2022). Both ascending and descending in social status necessitates cascades of molecular and genomic modifications to support the physiological and behavioral adaptations individuals need to acquire and maintain their new social status. An outstanding question is to what extent such molecular pathways are common to both social ascent and descent and which pathways are unique to gaining or losing status.

Experimentally, social ascent by subordinate males has been well-documented in Burton’s mouthbrooder cichlid fish, *Astatotilapia burtoni* (Huffman et al., 2012; Maruska & Fernald, 2013) and outbred CD-1 mice, *Mus musculus* (Williamson et al., 2017). In both species, ascending males rapidly decrease submissive behavior and increase aggression towards other males following removal of the alpha male. These dynamic behavioral changes are accompanied by a series of molecular and physiological changes. Within the brain, increases in immediate early gene (IEG) activity is observed during social ascent throughout the social decision-making network (SDMN), a system of evolutionarily conserved interconnected fore- and midbrain nuclei that strongly coordinate social behavior (O’Connell & Hofmann, 2011, 2012), as well as several other brain regions including the prefrontal cortex (Maruska, 2014; Williamson et al., 2019). Male *A. burtoni* cichlids also show increases in corticotropin-releasing factor (CRF) mRNA expression throughout the SDMN six hours after social ascent (Carpenter et al., 2014; Maruska et al., 2022). These changes are associated with an increase in HPG activity including increases in steroid hormone expression in the pituitary and receptor expression in the testes along with increases in circulating luteinizing hormone, follicle-stimulating hormone, androgens, estradiol, progesterone, and glucocorticoids in plasma. Other changes in body coloration, spermatogenesis and reproductive behavior occur from 6 to 24 hours following ascent (Huffman et al., 2012; Maruska, 2015). These modifications are followed by increases in androgen-ɑ and estrogen-β receptors in the SBN 120 hours post social ascent (Friesen & Hofmann, 2019; Maruska, 2014, 2015; Maruska & Fernald, 2013). Similarly, mice show increases in *GnRH* mRNA in the medial preoptic nucleus one hour following the onset of social ascent (Williamson et al., 2017).

Loss of social status has extraordinary effects on physiology and behavior, particularly for previously highly ranked individuals. For example, in mandrills, *Mandrillus sphinx* (Setchell & Wickings, 2005), and green anole lizards, *Anolis carolinensis* (Greenberg & Crews, 1990), alpha males who can no longer incur the physiological and energetic costs associated with maintaining dominance, can lose status dramatically leading to reduced mating opportunities and social isolation. Physiologically, this social descent has been associated with androgen-mediated reductions in body mass, testicular volume, sexual skin colorations, and scent gland activity (Greenberg & Crews, 1990; Setchell & Wickings, 2005). In cichlid fish, socially descending dominant males rapidly show submissive behavior, lose threatening facial and body markings, increase cortisol levels, reduce somatic growth, and exhibit distinct patterns of IEG activation across the SDMN distinct from ascending males (Hofmann et al., 1999; Maruska & Fernald, 2013). In mice, subordination experimentally induced through repeated social defeat by a larger conspecific has been shown to result in social avoidance, depressive and anxiety-like behavior as well as cognitive impairments (Wang et al., 2021). These changes are associated with numerous molecular and transcriptional changes that adapt the function of neural circuits particularly in the ventral tegmental area, nucleus accumbens, frontal cortex and hippocampus (Wang et al., 2021).

The medial amygdala (MeA) is a central hub of the SDMN, receiving input directly from olfactory processing areas and projecting outputs to several downstream nuclei that regulate behavioral output (Raam & Hong, 2021). As such, it is critical for the integration of sensory cues and internal states and regulates social behaviors including aggression and subordinate behaviors that are required for navigating a social hierarchy (Petrulis, 2020; Raam & Hong, 2021). Human imaging studies also show that activity in the amygdala is correlated with the rank of human faces in both stable and unstable hierarchies (Kumaran et al., 2016; Zink et al., 2008). In rhesus macaques, *Macaca mulatta*, increasing social status is associated with an increase in amygdala gray matter size (Noonan et al., 2014), lesions of the amygdala lead to individuals responding inappropriately to social cues and losing rank (Rosvold et al., 1954) and neuronal ensembles in the amygdala encode information about the social value of different ranks in a hierarchy (Munuera et al., 2018). The MeA is therefore a strong candidate region for encoding dominance status and regulating social hierarchy formation and reorganization.

In the present study, we developed a novel behavioral paradigm to concurrently investigate both social ascent and descent. Specifically, we took mice of equivalent social ranks from stable social hierarchies and placed them into new groups with unfamiliar animals of equivalent social rank. We predicted that male mice would form social hierarchies initially and would rapidly reestablish new and relatively stable hierarchies following social reorganization. We also predicted that animals that had previously been dominant would be faster to reestablish hierarchies than those animals that had previously been subordinate. Additionally, we tested whether individual attributes predicted an animal rank post social reorganization. We assayed plasma corticosterone levels before and at 70 minutes and 25 hours following social reorganization. We predicted that animals that were relatively more aggressive and had higher circulating corticosterone levels in their initial hierarchies would be more likely to attain higher rank following reorganization. We also predicted that animals of all ranks would show an increase in circulating corticosterone levels during social reorganization but that this would go back to baseline once hierarchies had stabilized. To examine the molecular modifications associated with social transitions, we profiled the MeA transcriptome 70 minutes following the onset of social ascent and descent to characterize common or differential gene expression patterns. We predicted that both social ascent and descent would lead to dramatically higher levels of gene expression of those genes that do not require *de novo* protein expression for their synthesis (Fowler et al., 2011) such as immediate early genes (IEGs) in the MeA compared to animals that did not change social rank. (as well as those regulating rapid neural modifications such as synaptic plasticity. We also predicted that a subset of genes and gene co-expression modules would be specifically associated with transitioning social status regardless of the direction of rank change.

## Results

### Male Mice form Social Hierarchies

All 65 groups of four male mice formed stable linear hierarchies with highly significant directional consistency (all p <0.001) prior to any social reorganization. The median and interquartile range of directional consistency of wins and losses across all groups was 0.98[0.92,1.00]. This indicates that all hierarchies were highly consistent with more dominant animals directing aggression towards more subordinate animals. Example raw sociomatrices of wins and losses are shown in **Figure 1B**. All raw sociomatrices are provided in **Supplemental Figures 2 & 3**.

**Figure 1:**
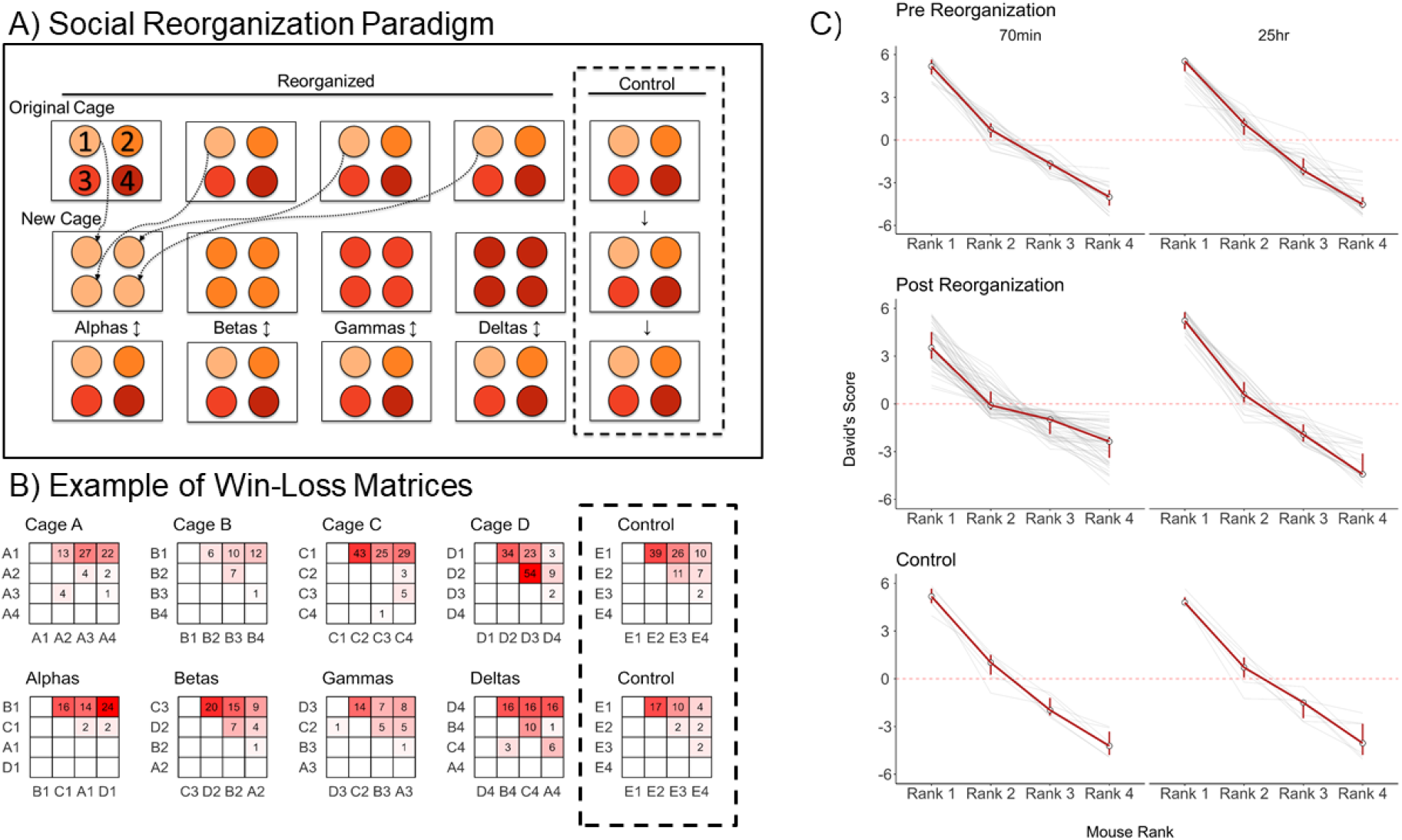
**A)** Illustration of social reorganization paradigm. Mice were placed in groups of 4 and subsequently reorganized by rank. Control animals remained in their existing group of four. **B)** Example win-loss matrices used to calculate David’s score for each animal. Values and degree of redness in cells represent the frequency of wins by animals in rows against animals in columns **C)** Red lines represent median David’s score with IQR of each social rank for control and reorganized animals pre- and post-reorganization. Gray lines represent individual animals and groups.

Each group had a distinct alpha male with a median and interquartile range of David’s score being 5.35[4.74,5.66] (**Figure 1C**). Beta males (rank 2) had David’s Scores that were typically above 0 (median[iqr] = 0.88[0.23,1.49]), indicating that, on average, beta males win more contests than they lose. Across these 65 groups, 54 beta males had positive David’s Scores, and only 11 had negative David’s scores. The majority of gamma (rank 3) males (64 out of 65) had David’s Scores below 0 (median[iqr] = -1.87[-2.42,-1.33]). Delta males (rank 4) were the most subordinate animal in each group, with the median and interquartile ranges of David’s score being -4.23[-4.74, -3.58]. As expected, there were no significant differences between groups that were in the reorganized and control conditions in the average David’s scores of each rank (One-Way ANOVA: all p>0.440, **Figure 1C**). Body mass at beginning of group housing was significantly associated with David’s Scores (**Supplemental Figure 4**, GLMM: β=0.19±0.06, p<0.005). However, this effect was very small. Indeed, many alpha and beta males were much smaller than delta and gamma males within individual hierarchies. Accordingly, body mass on day 7/8 during group housing was not significantly associated with David’s Scores (β=0.11±0.07, p<0.106).

**Figure 2:**
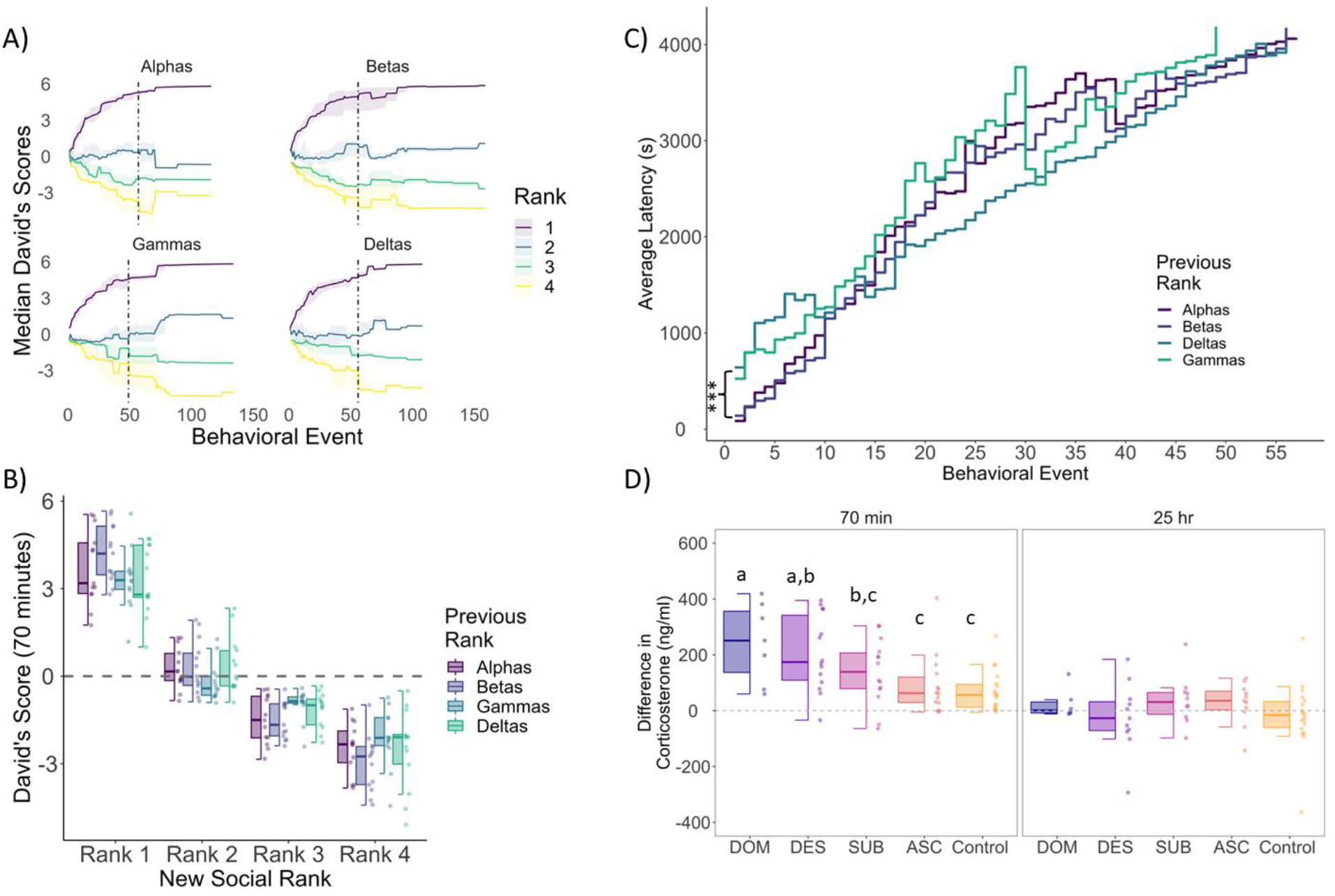
**A)** Emergence of social ranks post-social reorganization. Individual lines represent mean David’s score of animals by rank across each contest post-reorganization. **B)** Median and IQR of David’s score at 70 min post reorganization by rank. **C)** Average latency to engage in successive contests post-reorganization. Groups of previously alpha and beta males were significantly faster to engage in the first 8 agonistic behaviors compared to groups of previously gammas and deltas. **D)** Difference in plasma corticosterone from baseline (pre-reorganization) to 70 min and 25 hrs post-reorganization across social conditions. DOM = previously dominant males that remain dominant; DES = previously dominant males that socially descend; SUB = previously subordinate males that remain subordinate; ASC = previously subordinate males that socially ascend; Control = animals that were not socially reorganized

### Following social reorganization, male mice rapidly reestablish social hierarchies

Within 70 minutes of social reorganization, all 52 groups of equal-status male mice rapidly formed new linear social hierarchies. All groups showed highly significant directional consistency (p <0.001). The median and interquartile range of directional consistency of wins and losses across all groups was 1.00[0.89,1.00]. The emergence of individual ranks in new groups is shown in **Figure 2A**. For groups of all types (e.g. groups composed of all alphas, all betas, all gammas, all deltas), a new alpha male (purple line in **Figure 2A**) quickly rose to the top of the hierarchy. Over successive behavioral interactions, other animals determined their relative ranks. At 70 minutes, within all group types, all ranks had significantly different David’s Scores from each other (**Figure 2B**, post hoc tests: all p<0.020), confirming that new hierarchies emerged quickly.

We then asked whether animals with a prior history of winning were faster to engage in aggressive interactions than animals with a prior history of losing (**Figure 2C**). Groups of previously alpha males and groups of previously beta males were significantly faster to start fighting than cages of previously gamma (alpha vs gamma β=-1.36±0.35, p<0.001; beta vs gamma β=-1.06±0.33, p<0.005) or delta (alpha vs. delta β=-1.35±0.35, p<0.001; beta vs delta β=-1.05±0.35, p<0.005) males as determined by the latency to the first fight (**Figure 2C**). Previously alpha and beta males did not significantly differ in their latency to the first fight. Previously gamma and delta males also did not differ from each other in their latency to the first fight. Groups of previously alphas and betas continue to be significantly faster to fight than groups of previously gammas and deltas until the 9th behavioral event, when there are no significant differences in latency to fight between groups. This demonstrates that even animals that dramatically inhibited their aggression prior to social reorganization could engage rapidly in fighting when given the social opportunity. The total number of fights exhibited in the 70 minutes following social reorganization by groups of previously alpha, beta, gamma and delta males are shown in **Supplemental Figure 5.** Previously beta males had significantly more contests on average over the 70 minute period than all other ranks (vs alphas β=0.26±0.07, p<0.001; vs gammas β=0.37±0.07, p<0.001; vs deltas β=0.27±0.07, p<0.001), whereas previously alpha, gamma and delta groups did not significantly differ from each other.

### Aggression received prior to reorganization predicts social ascent

We examined whether it was possible to predict which animals rose to the top or fell to the bottom of their new hierarchies post-reorganization. We found that neither body mass nor David’s Score in the pre-reorganization group significantly predicted David’s Score in the new hierarchy within groups of all alphas, all betas, all gammas or all deltas. Raw rates of aggression exhibited before reorganization also did not predict David’s Score post reorganization for each rank. However, we did find that rates of aggression received prior to reorganization were positively associated with later David’s Scores in groups of beta (β=0.69±0.26, p=0.014), gamma (β=0.29±0.20 p=0.144) and delta (β=0.40±0.22, p=0.080) males (**Supplemental Figure 6A**). Although this effect was only significant among previously beta males, this suggests that among lower ranked individuals, individuals who received relatively more aggression before the reorganization were more likely to socially ascend after hierarchy reorganization.

We also tested whether animals that were housed together before reorganization showed similar propensities to socially ascend or descend following reorganization. For all 52 pre-reorganization groups that were reorganized, we calculated the sum of the differences in their post-reorganization ranks. We found that the sum of all these differences was significantly lower than a theoretical distribution of differences from random samples (**Supplemental Figure 6C**, p<0.001). This group-of-origin effect indicates that social dynamics prior to reorganization predisposed individuals living together to be more likely to attain similar ranks following their reorganization. Total aggression occurring within each group prior to reorganization may be reflective of such group dynamics. For groups comprised of betas (β=0.20±0.10, p=0.047), gammas (β=0.17±0.08, p=0.053) and deltas (β=0.13±0.09, p=0.151), we found that higher rates of aggression per group prior to reorganization were associated with an increased likelihood of becoming dominant, although this effect was only significant in previously beta males (**Supplemental Figure 6B**).

### Reestablished Social Hierarchies Remain Stable over 25 hours

Observations of disrupted hierarchies (N=24 groups; 6 groups/social condition) were recorded up to 24hrs after reorganization to examine whether hierarchies remained stable. Regardless of social condition, all animals that attained rank 1 (alpha males) after 70 minutes continued to possess the highest David’s Score on day 2 indicating that the alpha position was highly robust. Indeed, none of these rank 1 animals lost to any other animal on day 2 (Binomial test: 0/24, p < 0.001). Of the 24 males ranked 2 after day one, only 3 lost more fights to a lower-ranked male than they won on day 2 (Binomial test: 3/24, p < 0.001). Two of these rank 2 males came from previously gamma and delta conditions, where they lost only one more fight than they won against a lower-ranking male. Of the 24 animals ranked 3 after day one, only 1 from the previously gamma social condition lost one fight more than they won against a lower-ranking animal (Binomial test: 1/24, p < 0.001). These data demonstrate the remarkably high consistency of ranks across the first day of housing together. This is also observable from the consistency of David’s Scores across 25 hours (**Figure 2A**).

### Social rank is not associated with corticosterone levels in stable hierarchies

We tested whether social status was associated with basal circulating levels of corticosterone. Corticosterone levels were assessed on all groups of four male mice in stable hierarchies one week after group housing began, three days prior to the social reorganization. As expected, we found no significant differences between the plasma corticosterone levels of animals who were assigned to the control 74.7ng/ml [53.9, 100.0] or reorganized 88.6ng/ml [63.7, 124.0] conditions (**Supplemental Figure 7A**). Individual social rank also had no overall effect on plasma corticosterone levels (**Supplemental Figure 7B**; alphas 87.3 ng/ml [71.0,117], gammas 71.6 ng/ml [55.3,108], and deltas 82.4 [53.4,128]).

### Previously dominant animals exhibit the greatest elevation in corticosterone following social reorganization

We examined whether circulating corticosterone levels reflected an animal’s social status in response to social reorganization. We found that all groups had higher plasma corticosterone levels 70 minutes post-reorganization compared to pre-reorganization (Ascenders t_13_=3.2, p=0.007; Subordinates t_13_=4.5, p<0.001; Descenders t_13_=5.7, p<0.001; Dominants t_6_=4.6, p=0.003) (**Figure 2D**). Notably, controls, which were placed into a new cage to be consistent with all other groups, had higher corticosterone even though they were not socially reorganized (Control t_17_=4.1, p<0.001). We found that animals that were previously alpha males showed larger increases in corticosterone during reorganization compared to controls, regardless of whether they maintained their status (dominants - β_17_=7.13 ± 48.87, p<0.001) or lost their status (descenders - β_1_=38.10 ± 39.67, p<0.001). Previously subordinate animals did not show significant elevations in corticosterone compared to controls, although subordinates who remained subordinate did show a non-significant increase (subordinates - β=67.99 ± 38.69, p=0.085). No significant differences in corticosterone increases were observed between previously dominant animals that remained dominants (dominants) or lost status (descenders) or between previously subordinate animals that remained subordinates (subordinates) or gained status (ascenders). At 25 hours after social reorganization, we observed no differences in plasma corticosterone compared to pre-reorganization levels for any group. Further, there were no significant differences in corticosterone between groups, demonstrating the rapidity with which hierarchies stabilize following social reorganization (**Figure 2D**).

### Social hierarchy reorganization causes substantial changes in medial amygdala transcriptome profiles of male mice

Given its critical role in regulating the coordination of aggressive and subordinate behavior in social hierarchies, we examined the MeA transcriptomes of ascending and descending males in comparison to dominant and subordinate animals who did not change rank. The total number of differentially expressed genes (DEGs) for each comparison group are given in **Table 1** and **Supplemental Figures 8 & 9**.

**Table 1.** Total DEGs between all 6 comparisons split across four different log2 fold change thresholds. DOM = previously dominant males that remain dominant; DES = previously dominant males that socially descend; CDOM = control dominant animals that remain dominant. SUB = previously subordinate males that remain subordinate; ASC = previously subordinate males that socially ascend; CSUB = control subordinate animals that remain subordinate.

|  | *Previously Dominant* |  |  |  |  |  | *Previously Subordinate* |  |  |  |  |  |
| --- | --- | --- | --- | --- | --- | --- | --- | --- | --- | --- | --- | --- |
|  | DES vs. DOM |  | DES vs. CDOM |  | DOM vs. CDOM |  | ASC vs. SUB |  | ASC vs. CSUB |  | SUB vs. CSUB |  |
| Log Fold Change | ↑↑ | ↓↓ | ↑↑ | ↓↓ | ↑↑ | ↓↓ | ↑↑ | ↓↓ | ↑↑ | ↓↓ | ↑↑ | ↓↓ |
| ≥ 0.20 | 1187 | 1130 | 575 | 526 | 802 | 832 | 658 | 727 | 280 | 225 | 309 | 322 |
| 0.20 – 0.75 | 990 | 888 | 463 | 472 | 676 | 719 | 576 | 656 | 250 | 185 | 276 | 248 |
| 0.75 – 1.50 | 176 | 212 | 100 | 51 | 107 | 104 | 77 | 61 | 26 | 37 | 36 | 62 |
| > 1.50 | 21 | 30 | 12 | 3 | 19 | 9 | 5 | 10 | 4 | 3 | 6 | 12 |

The largest differences in gene expression were found when comparing animals that changed social status to those animals that were also socially reorganized but maintained their social status. Notably, when considering log_2_ fold-changes > 0.2, animals that were previously dominant and socially descended (DES) had 2,317 (15.3% of all genes in our analysis) differentially expressed genes compared to other alpha males that maintained their dominance (DOM) in reorganized groups, whereas animals that were previously socially subordinate and socially ascended (ASC) had 1,385 genes ( 9.12%) that were differentially expressed compared to subordinates that remained subordinate in reorganized groups (SUB). This reflects 1.7 times more genes being differentially expressed in socially descending animals than in socially ascending animals. When considering log_2_ fold-changes greater than 1.5, socially descending animals had 3.3 times more differentially expressed genes than socially ascending animals when compared to animals that were reorganized but did not change ranks (51 genes in descenders vs. 15 genes in ascenders) (**Table 1**). When comparing reorganized dominant animals to control dominants (CDOM), those reorganized animals that maintained their dominance status had around 33% more differentially expressed genes compared to those that socially descended (1,634 vs. 1,101 DEGs). This was also approximately two times the number of DEGs identified when comparing reorganized subordinate animals who either maintained rank or socially ascended to control subordinates, further demonstrating that social reorganization leads to greater transcriptomic changes in dominant compared to subordinate males. (**Table 1**).

We performed GO-enrichment analysis on differentially expressed genes for both dominant and subordinate animals (**Supplemental Figures 8 & 9**). Most strikingly, we observed that socially descending animals had an over-representation of DEGs related to myelination, axon ensheathment, gliogenesis, glial cell, and oligodendrocytes differentiation. Animals descending in status had absolute log_2_ fold reductions of many genes related to these processes above 0.75-fold change when compared to both animals maintaining dominant status and control dominants. These genes include myelin oligodendrocyte glycoprotein (*Mog*), lysophosphatidic acid receptor 1 (*Lpar1*), myelin and lymphocyte protein (*Mal*), myelin basic protein (*Mbp*), myelin associated oligodendrocyte basic protein (*Mobp*), cyclic nucleotide phosphodiesterase (*Cnp*), tetraspanin-2 (*Tspan2*), NK6 homeobox 2 (*Nkx6-2*), contactin 2 (*Cntn2*), rho guanine nucleotide exchange factor 10 (*Arhgef10*), brain enriched myelin associated protein 1 *(Bcas1*), myelin associated glycoprotein (*Mag*), myelin regulatory factor (*Myrf*), proteolipid protein 1 (*Plp1*), and oligodendrocytic myelin paranodal and inner loop protein (*Opalin*). Moreover, several of these myelination genes were significantly more highly expressed in animals maintaining dominant status after reorganization than control dominants (e.g. *Mal, Mag, Mog, Mobp, Cnp, Bcas1, and Lpar1),* suggesting that not only are descending animals downregulating these processes, but animals maintaining dominant status are upregulating them (**Figure 3A**).

**Figure 3.**
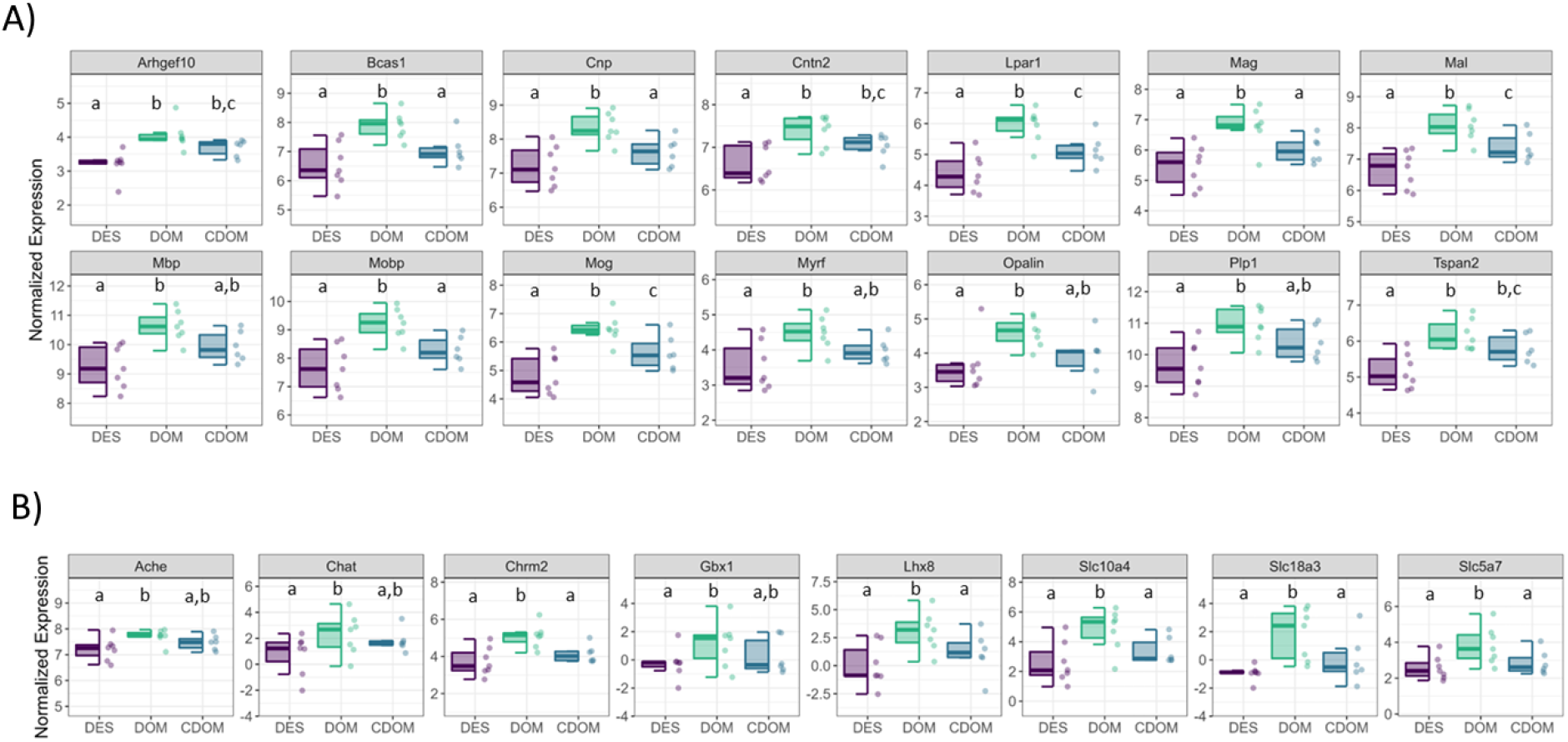
Log normalized counts of most differentially expressed **A)** myelination genes, and **B)** cholinergic signaling genes in dominant animals. DOM = previously dominant males that remain dominant; DES = previously dominant males that socially descend; CDOM = control dominant animals that remain dominant.

Compared to socially descending animals, the MeA of males that maintained dominant status in reorganized hierarchies had an over-representation of differentially expressed genes associated with cholinergic signaling. Indeed, 7 out of the top 25 most differentially expressed genes in the DES vs DOM comparison were such genes. In particular, LIM homeobox 8 (*Lhx8*, fc=3.764), solute carrier family 18 member A3 (*Slc18a3*, fc=3.07), solute carrier family 10 member 4 (*Slc10a4*, fc=2.55), solute carrier family 5 member 7 (*Slc5a7*, fc=1.56), choline O-acetyltransferase (*Chat*, fc=1.67), cholinergic receptor muscarinic 2 (*Chrm2*, fc=1.59), gastrulation and brain-specific homeobox protein 1 (*Gbx1*, fc=1.75), and acetylcholinesterase (*Ache*, fc=0.506) are all relatively more highly expressed in maintaining dominants than dominants that socially descend (**Figure 3B**). Several of these genes (*Lhx8* fc=2.37; *Slc18a3* fc=2.09; *Slc10a4* fc=1.44; *Slc5a7* fc=1.17; *Chrm2* fc=1.02) are also significantly more highly expressed in the DOM vs CDOM comparison. Reorganized subordinate animals do not show any substantial changes in expression of cholinergic related genes, although those individuals maintaining subordinate status did show relatively higher expression of *Ache* (fc=0.52), *Chrm2* (fc=1.03), and *Chrnb2* (fc=0.22) compared to socially ascending animals.

Finally, we examined gene expression patterns of IEGs between each comparison. In support of our prediction that social transition would activate IEGs, we observed that significantly more of these genes showed relatively higher expression in socially descending animals (23.3% of IEGs) compared to animals that stay dominant (6.9%) (Binomial Test: *p=0.002*) (**Supplemental Figure 10**). The *genes Bdnf, Cort, Egr2, Egr3, Egr4, Nr4a2, Fosb*, and *Junb* were some of the most highly differentially expressed in socially descending animals. However, there were many fewer IEGs differentially expressed between socially ascending animals (5.2%) and those that stayed subordinate (8.6%).

### WGCNA reveals co-expressed gene modules uniquely associated with social descent and ascent

We used WGCNA of all samples to infer gene networks associated with change in social status. This analysis revealed 25 gene co-expression modules (**Figure 4A**). Three modules (darkturquoise, darkgreen, pink) had significantly differentially expressed module eigengenes (deMEGs) when comparing descending animals to both dominant groups (DOM and CDOM). Five other modules (salmon, brown, green, black, yellow) had significant deMEGs when comparing descending animals to one of the other dominant groups. **Figure 4B** shows module eigengene expression across each social condition and summarizes hub genes (i.e., genes with a module membership (MM) > 0.8) and GO terms significantly associated with each module.

Descending animals had greatly reduced module eigengene (ME) expression compared to rearranged dominants that maintained status in the yellow module (DES vs DOM β=1.72 ± 0.38 *p<0.001*). GO-enrichment analysis revealed that these genes were largely involved in central nervous system myelination, axon ensheathment, oligodendrocyte development. Genes related to myelination production by oligodendroyctes with MM > 0.8 included several of those identified by our DEG analysis above, including *Tspan2, Cnp*, *Plp1*, *Mobp*, *Mbp*, *Mog*, *Mag*, *Mal*, *Bcas1*, *Lpar1*, *Cntn2* (**Figure 3A**). Descending animals had significantly increased deMEG expression compared to both dominant groups in the darkturquoise module which consists of only 64 genes (DES vs DOM β=-1.55 ± 0.40 *p<0.001*; DES vs CDOM β=-1.22 ± 0.42 *p<0.001*). GO-enrichment analysis suggested that these genes were related to regulation of synapse organization and postsynaptic activity. Significantly differentially expressed genes included *Cacnb1* (voltage-dependent L-type calcium channel subunit beta-1, fc=0.41), *Cnih2* (protein cornichon homolog 2, fc=0.37), *Thop1* (thimet oligopeptidase, fc=0.59), *Lmtk3* (lemur tyrosine kinase 3, fc=0.50) and *Dlgap3* (discs large homolog associated protein 3, fc=0.36). Interestingly, both descending and dominant animals who experienced the disruption of social reorganization had significantly increased deMEG in the salmon module. The salmon module consists of 238 total genes. GO-enrichment analysis indicated that these genes are involved in sugar and carbohydrate metabolic processes.

Fewer modules were differentially expressed between ascending and subordinate groups (**Figure 4**). Indeed, no module showed consistent differences between ascenders and both subordinate conditions. However, we did find that animals that maintained their subordinate status after reorganization had consistently higher deMEG expression compared to both ascenders and control subordinates in the orange module (SUB vs ASC β=1.18 ± 0.42 *p=0.013*; *SUB* vs CSUB β=1.46 ± 0.44 *p=0.004*). GO-enrichment analysis indicated that genes in this module are related to mononuclear cell and lymphocyte differentiation. Strikingly, both ascending and descending animals showed reduced ME gene expression in the pink module compared to controls (**Figure 4B**). Based on GO-enrichment analysis, genes in this module appear related to nervous system development, neurogenesis, synapse organization and negative regulation of kinase activity.

**Figure 4.**
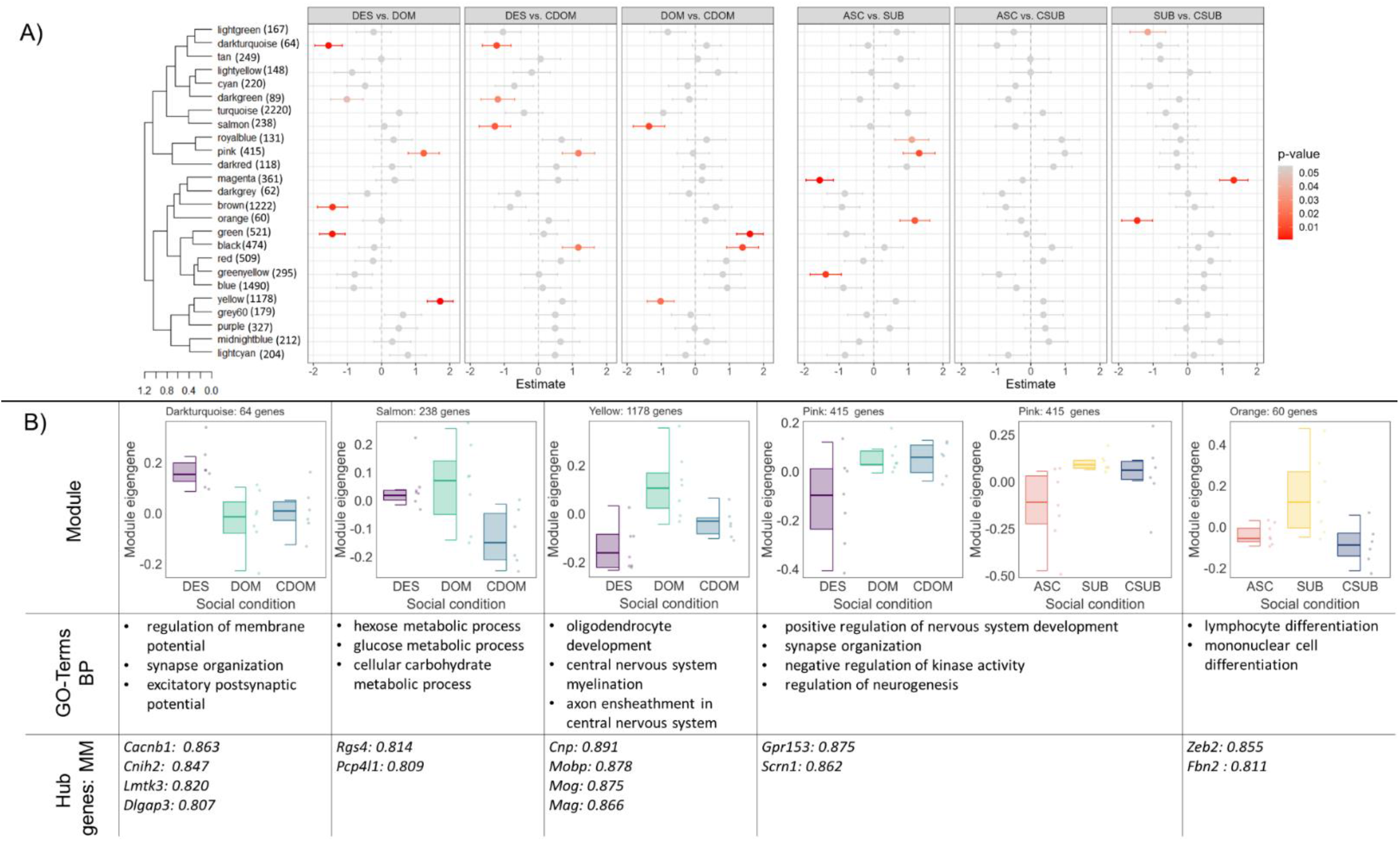
WGCNA module analysis. **A)** Linear regression estimates of eigengene expression of WGCNA modules for each social comparison. Module sizes are noted next to each module. **B)** Boxplots represent module eigengene expression by social condition and points represent individual subjects. A summary of biological processes GO-terms and genes with high module membership (MM) are noted for each module. DOM = previously dominant males that remain dominant; DES = previously dominant males that socially descend; CDOM = control dominant animals that remain dominant. SUB = previously subordinate males that remain subordinate; ASC = previously subordinate males that socially ascend; CSUB = control subordinate animals that remain subordinate.

### Social descent and social ascent share transcriptomic signatures in the MeA

Finally, we asked whether we could identify genes associated with social transition independent of the direction of status change, as such genes might reveal molecular pathways involved in the plastic responses required to adjust brain and behavior to a major life history transition more generally. To do this, we asked whether there was concordance in expression variation between that were differentially expressed in descenders versus previously dominant animals who maintained status (DES vs. DOM) on the one hand and ascenders versus subordinates that remained subordinate (ASC vs. SUB) on the other. We found that the descenders and ascenders shared 279 DEGs out of the 3144 unique genes that were differentially expressed in either comparison (**Figure 5, Supplemental Table 1**). Remarkably, 265 out of these 279 DEGs were differentially expressed in the same direction, with 142 being more highly expressed in animals changing social status and 123 showing reduced expression in animals changing social status. Only 14 out of 279 genes showed opposite patterns of expression change in individuals changing social status. This consistency in the direction of gene expression changes between ascending and descending males was highly significant (Chi-Squared test: χ2=222.03, p<0.001, Φ=0.89).

**Figure 5.**
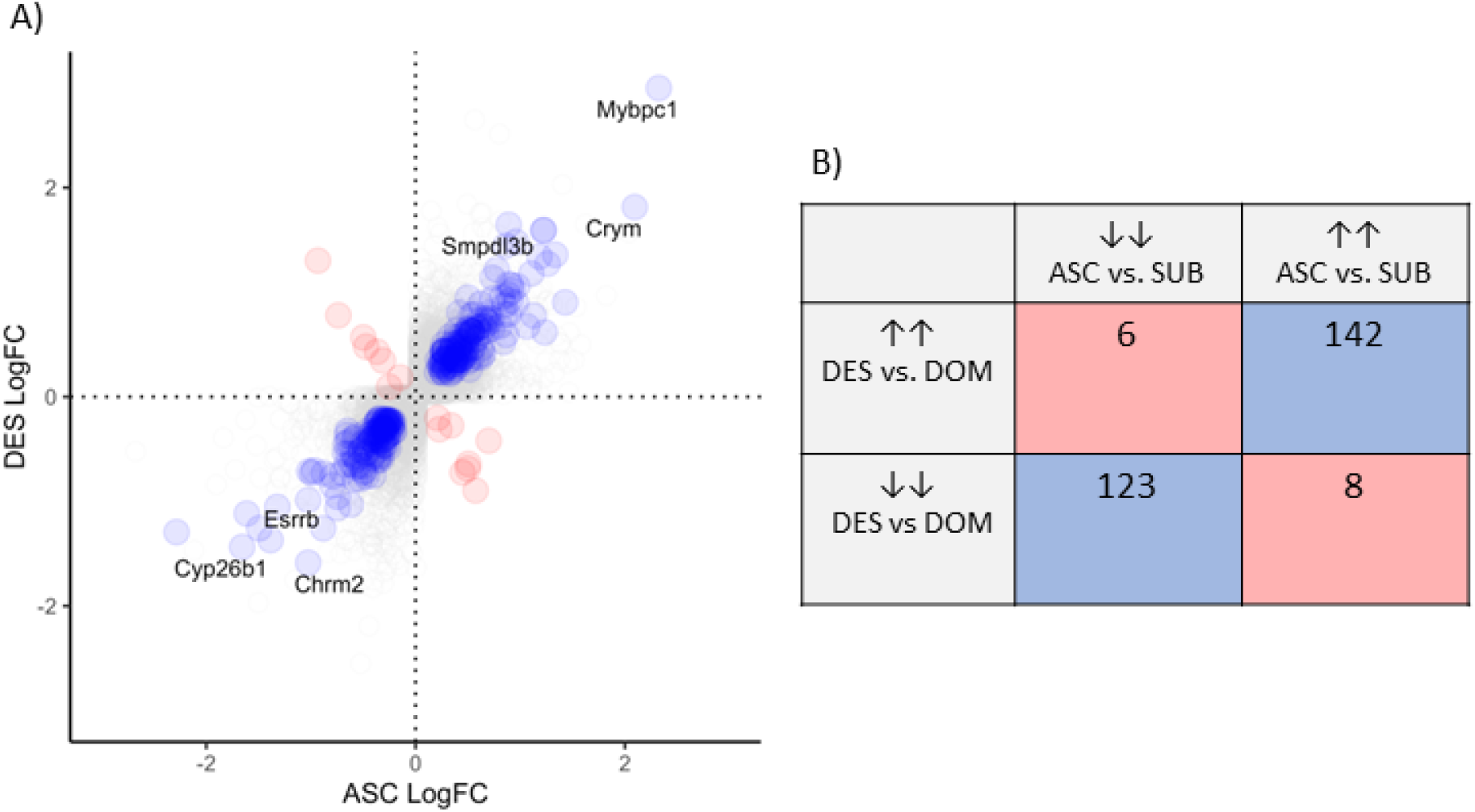
Concordance of DEGs in social ascenders and descenders compared to animals who maintained their social status. **A)** Relative fold change in ASC vs SUB (x-axis) against relative fold change in DES vs DOM (y-axis). Positive values represent relatively higher expression in animals transitioning social status (ASC and DES). Blue dots represent genes showing congruence in fold changes across both comparisons. Red dots genes showing significant changes in opposite directions. **B)** Number of DEGs showing significant differences in expression in each comparison. DOM = previously dominant males that remain dominant; DES = previously dominant males that socially descend; SUB = previously subordinate males that remain subordinate; ASC = previously subordinate males that socially ascend.

The gene with the largest fold change in both comparisons was myosin binding protein C1 (*Mybpc1*; DES vs DOM: fc = 2.95; ASC vs SUB fc=2.32, **Figure 5A**). This gene is a molecular marker for a subset of vasoactive intestinal polypeptide-expressing (VIP) interneurons (Jiang et al., 2023). Notably, *Vip* (fc=1.77) was the seventh most differentially expressed gene in the DES vs DOM comparison showing relatively higher expression in descenders. Conversely, VIP receptor 1 (*Vipr1*, fc=0.87) was relatively more highly expressed in ascenders compared to subordinates. Another gene that showed remarkably high relative expression in transitioning animals was μ-crystallin (*Crym*), a thyroid hormone binding protein. This was the second most differentially expressed gene in the ASC vs SUB comparison (fc=2.09), and the sixth most differentially expressed in the DES vs DOM comparison (fc=1.81). Several other genes related to thyroid hormone signaling were also differentially expressed (**Supplemental Figure 11**). Descending males had significantly higher expression of thyroid hormone receptors a (*Thra*, fc=0.29) and b (*Thrb*, fc=0.58), thyrotropin-releasing hormone receptor (*Trhr*, fc=0.90) and thyroid releasing hormone (*Trh*, fc=1.75). Ascending males also had significantly higher expression of *Trhr* (fc=0.96) compared to reorganized males that remained subordinate. Further, reorganized dominants that descended or stayed in the same rank had significantly higher expression of iodothyronine deiodinase 2 (*Dio2*) compared to control dominants (DES vs. DOM: fc=1.22, DES vs. CDOM: fc=0.96) *Dio2* encodes an enzyme that converts the prohormone thyroxine (T4) to the active thyroid hormone 3,3’,5-triiodothyronine (T3). Descenders also had significantly higher levels than control dominants of iodothyronine deiodinase 3 (*Dio3*, fc=1.86), which encodes an enzyme that inactivates the production of circulating thyroid hormone.

To more stringently characterize genes associated with social transition, we further examined which genes showed differential expression in both descenders compared to dominants and control dominants and in ascenders compared to subordinates and control subordinates. This comparison revealed far fewer genes, with five genes (*Car12, Unc13a, Lpl, Tesc, Pitpnm2*) having higher expression in both ascenders and descenders compared to other groups, and three genes (*Rai2, Kif21a, Nap1l5*) showing lower expression in both ascenders and descenders compared to other groups (**Supplemental Figure 12**). No genes showed discordant expression patterns between ascenders and descenders, indicating that social transitions are significantly associated with concordant patterns of gene expression independent of the direction of the social status change (Chi-Squared test: χ2=4.30, p=0.038, Φ=0.73).

To further identify common transcriptomic profiles between animals that socially ascend and descend, we applied WGCNA only to socially reorganized animals. We compared socially transitioning animals (ascenders and descenders = TRN) to animals that remained subordinate (SUB) or remained dominant (DOM). WGCNA identified 20 modules, of which 8 showed significantly different ME expression between transitioning animals and at both non-transitioning groups (**Figure 6**).

**Figure 6.**
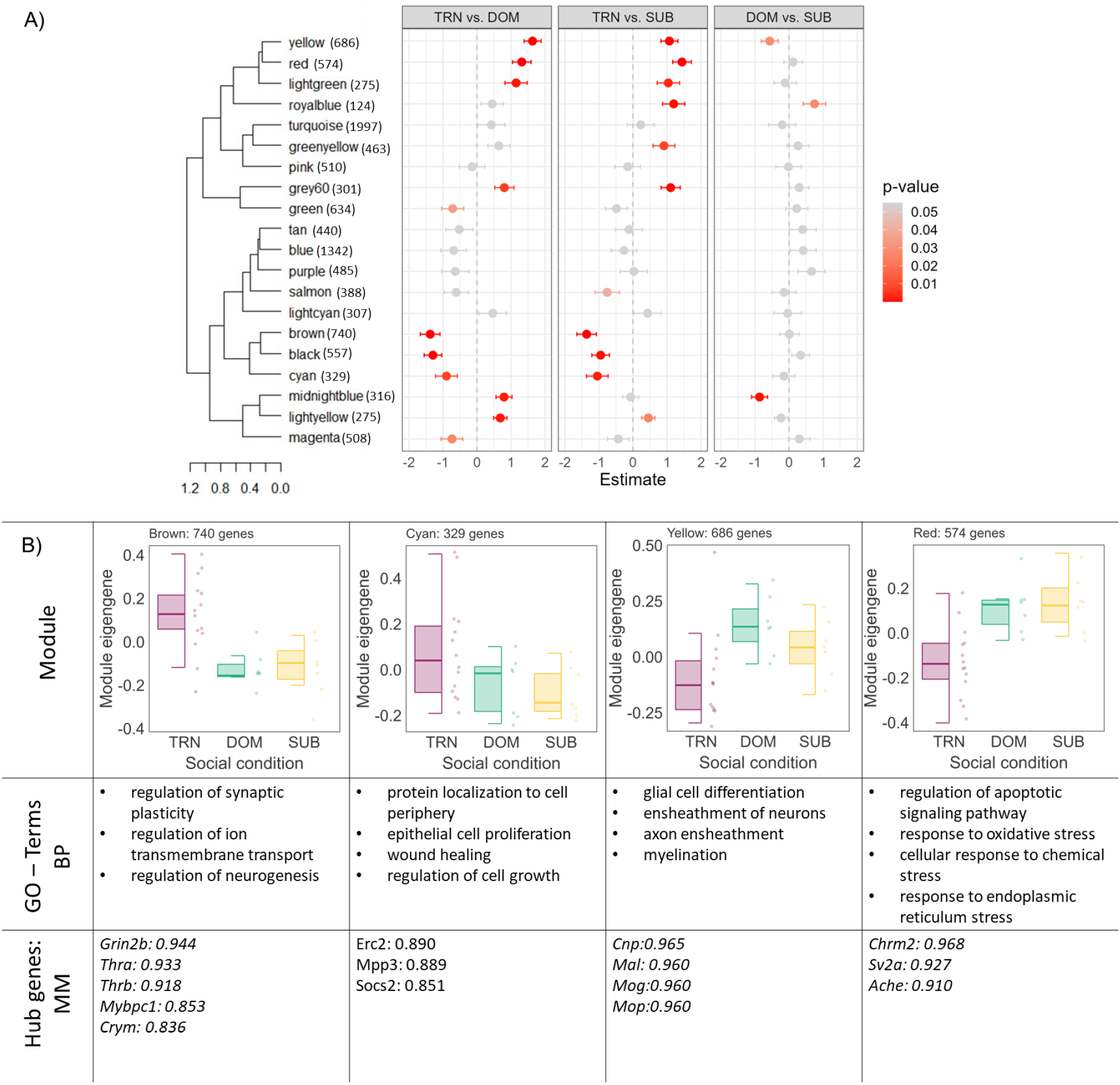
Modules identified by Weighted Gene Correlation Network Analysis (WGCNA) of social transition. Module sizes are noted next to each module. **A)** Linear regression estimates of eigengene expression of WGCNA modules against social conditions. **B)** Boxplots represent module eigengene expression by social condition, and points represent individual subjects. For each module, biological processes, GO-terms, and genes with high module membership are noted. TRN = previously dominant or subordinate animals that transitioned rank; DOM = previously dominant males that remain dominant; SUB = previously subordinate males that remain subordinate.

Three modules (brown TRN vs DOM β=-1.37 ± 0.28 p<0.001, TRN vs SUB β=-1.36 ± 0.28 p<0.001; black TRN vs DOM β=-1.29 ± 0.25 p<0.001, TRN vs SUB β=-0.95 ± 0.25 p<0.001; and cyan TRN vs DOM β=-0.89 ± 0.31 p=0.007, TRN vs SUB β=-1.04±0.31 p=0.002) had higher deMEG expression in transitioning animals compared to both dominants and subordinates (**Figure 6**). GO-enrichment indicated that genes belonging to the brown module are involved in the regulation of neurogenesis, synaptic plasticity, and membrane potential. 83 out of the 92 hub genes (90.2%) were differentially expressed in at least one comparison, with 43 (46.7%) being differentially expressed in both the DES vs DOM and ASC vs SUB comparisons. The hub gene with the highest MM is Glutamate Ionotropic Receptor NMDA Type Subunit 2B (*Grin2b*, MM=0.94), which is differentially expressed in both comparisons (fc_des-dom_ =0.52; fc_asc-sub_ =0.43). Notably, *Crym* (MM=0.84) and *Mypb1c* (MM=0.85) are also hub genes of this module. Genes in the cyan module are related to processing of ncRNAs and rRNAs, as well as rRNA and tRNA metabolic and mRNA catabolic processes. Two genes, NMDA receptor synaptonuclear signaling and neuronal migration factor (*Nsmf*, fc_des-dom_ == 0.40, fc_asc-sub_ = 0.47) and ELKS/RAB6-interacting/CAST family member 2 (*Erc2* fcd_es-dom_ = 0.37, fc_asc-sub_ = 0.46) are significantly differentially expressed in both comparisons. GO-enrichment suggested that the black module contained genes related to histone modification, methylation, and peptidyl-lysine modification. Notably, 64 out of the 388 genes of this module showed significantly higher expression in descenders compared to dominants, but only 1 gene showed significantly higher expression in ascenders compared to subordinates.

Five modules (yellow TRN vs DOM β=1.63 ± 0.25 p<0.001, TRN vs SUB β=1.07 ± 0.25 p<0.001; red TRN vs DOM β=1.31 ± 0.27 p<0.001, TRN vs SUB β=1.44 ± 0.27 p<0.001; lightyellow TRN vs DOM β=0.68 ± 0.2 p<0.001, TRN vs SUB β=0.45 ± 0.2 p=0.020; lightgreen TRN vs DOM β=1.15 ± 0.33 p<0.001, TRN vs SUB β=1.04 ± 0.33 p=0.003; and grey60 TRN vs DOM β=0.81 ± 0.28 p=0.003, TRN vs SUB β=1.12±0.28 p<0.001) had lower deMEG expression in transitioning animals compared to both dominants and subordinates (**Figure 6**). Go-enrichment analysis of the yellow module indicated myelination and axon ensheathment as biological processes associated with these genes. Of the 686 genes in this module, 110 were hub genes with 104 (95%) being differentially expressed in descenders compared to dominants. These include myelination genes: *Cnp, Bcas1, Mal, Mog, Mbp, Mob*, and *Lapr*. However, only one of the hub genes, radixin (*Rdx*) also showed differential expression in ascenders compared to subordinates (fc_des-dom_ = 0.50; fc_asc-sub_ = 0.31). In the red module, 73 genes out of 574 were identified as hub genes with 66 of these being differently expressed in at least one of the comparison groups. Two hub genes with MM > 0.9 of particular interest are those involved in cholinergic signaling, *Chrm2* (fc_des-dom_ = -1.59, fc_asc-sub_ =-1.03) and *Ache* (fc_des-dom_ = 0.51; fc_asc-sub_ = 0.52), which are relatively less expressed in both ascending and descending animals.

## Discussion

In the present study, we demonstrated that CD-1 male mice can rapidly reorganize themselves into linear dominance hierarchies following a social challenge leading mice to socially descend, ascend or remain the same rank in their new hierarchies. Previously dominant animals showed the largest response to this social reorganization, exhibiting significantly higher levels of plasma corticosterone and a two-fold increase in DEGs in the MeA. Additionally, we identify a subset of genes involved in the regulation of synaptic plasticity and likely related to learning and memory processes that are commonly differentially expressed in mice who socially transitioned in either direction (ascent or descent).

### Animals rapidly reestablish social hierarchies

We observed that all groups of four male outbred CD-1 mice rapidly formed social hierarchies. Further, when animals were socially reorganized and placed into new groups with novel conspecifics of equivalent rank, new hierarchies were established within minutes. These new hierarchies were stable for at least 25 hours. Groups composed of formerly all alpha or all beta males were the quickest to engage in dominance behavior, but groups of all gamma and all delta animals also very rapidly began to determine social ranks. The readiness of low-ranked individuals to social ascend when given the opportunity indicates that social subordination is a product of the social environment and not a default phenotype of individuals. We have previously demonstrated that beta CD-1 males will rapidly rise to dominant status following the removal of alpha males (Williamson et al., 2017; Williamson et al., 2019), but this is the first demonstration that mice of all social ranks rapidly adjust their aggression and subordinate behavior following changes in their social environment. Our findings are also broadly consistent with studies in Burton’s mouthbrooder cichlid, *A. burtoni* (Friesen et al., 2022; Chase et al., 2002; Maruska, 2015), monk parakeets, *Myiopsitta monachus* (Hobson & DeDeo, 2015), and rhesus macaques, *M. mulatta*, (Snyder-Mackler et al., 2016), which all demonstrate that animals will readily reorganize themselves into stable social hierarchies when placed into novel social groups. These findings also support the hypothesis that dominance hierarchies are a universal feature of social groups and emerge to maintain social equilibrium (Chase et al., 2022).

Within our social reorganization paradigm, we were able to identify animals that maintained their social rank, lost social rank, or gained social rank. Therefore, within the same study we can investigate both social ascent and social descent. This is a unique feature of our study design as most commonly studies focus solely on either social defeat or social ascent (Maruska, 2015; Wang et al., 2021; Williamson et al., 2016). Notably, neither body mass nor previous dominance level (David’s Scores) within the initial hierarchy predicted which animals were likely to socially ascend or descend. However, we did find that individuals who received relatively more aggression before the reorganization were more likely to socially ascend. This suggests that those animals who are willing to engage in agonistic contests, even if they lose most of the time, are more likely to seize social opportunities successfully when they emerge. Similarly, we found that previously beta males were the animals who exhibited the highest levels of aggression following reorganization, indicating that animals who have experienced high amounts of both winning and losing are willing to continue to fight for dominance status for longer. We also found that animals from the same hierarchy prior to reorganization were more likely to show similar ranks in their new hierarchies post-reorganization. This suggests that some aspect of the social dynamics of initial hierarchies, likely the levels of aggression received by animals from dominant males, has long-term effects on the social dominance behavior of individuals.

### Dominant males exhibit the largest changes in corticosterone and transcription profiles following social reorganization

When we assessed the impact of social reorganization on changes in plasma corticosterone levels, we found that previously dominant (alpha) males had the highest increases in plasma corticosterone 70 minutes following the social reorganization compared to previously subordinate (gamma or delta) males. This was true for previously dominant animals that maintained their status, or those that socially descended. Indeed, subordinates did not show higher levels of corticosterone compared to control animals who were not socially reorganized. This suggests that dominant males may find social reorganization more physiologically stressful than subordinate individuals. Notably, the plasma corticosterone levels of all animals returned to baseline 25 hours following the social reorganization, suggesting that any effects of this social stress may be highly transient. Although we did not find any differences in corticosterone levels between dominant and subordinate animals at baseline, this is not unexpected (Williamson, Lee, et al., 2017). Indeed, our findings are broadly consistent with studies that have demonstrated that across vertebrates plasma glucocortocoid levels are higher in dominant animals during periods of highly competitive interactions between dominant animals such as during high reproductive competition or food shortage (Beehner & Bergman, 2017; Cavigelli & Caruso, 2015; Fox et al., 1997; Maguire et al., 2021; Milewski et al., 2022; Sapolsky, 1993; Wingfield et al., 1990).

Social reorganization induced more transcriptional variation in dominant animals versus subordinate animals. When compared to non-reorganized control animals of equivalent ranks, reorganized dominants had approximately twice as many DEGs as did subordinate animals. When considering fold changes greater than 0.2, we also found that reorganized dominant animals had 1.7 times higher number of genes with differential expression between those that maintained or lost status when compared to reorganized subordinate animals that either maintained or gained status. When considering genes with large fold changes (>1.5), dominants vs descenders had 3.3 times as many differentially expressed genes as subordinates vs ascenders (51 vs 15 DEGs). In agreement with this result, we found that a significantly higher proportion of IEGs such as *Bdnf*, *Egr2*, *Egr3*, *Fosb* and *Junb* were relatively more highly expressed in descending animals. We also identified a WGNCA module (darkturquoise) containing genes related to the regulation of synapse organization and postsynaptic activity that were relatively more highly expressed in descending animals. These findings suggest that the effects of social reorganization on changes in MeA gene expression appear to be exacerbated in dominant animals and particularly those that lose social status. Our data support evidence demonstrating that this region is particularly sensitive to social challenges with experience of aggression and social defeat in mice leading to large transcriptional changes (Cordner et al., 2021; Saul et al., 2017; Yamaguchi et al., 2020). Further, our findings also support a role for transcriptional remodeling of the MeA occurring during social status transitions as has been suggested by early immediate gene studies of social ascending and descending *A. burtoni* cichlids (Maruska & Fernald, 2013) as well as socially defeated rats, *Rattus norvegicus* (Fekete et al., 2009) and Syrian hamsters, *Mesocricetus auratus* (Morrison et al., 2014).

DEG and WGCNA analysis confirmed that multiple genes related to myelination and oligodendrocyte differentiation had greatly elevated expression in dominants that maintained their status compared to descending dominant animals. These genes include those affect myelin structure and composition such as myelin basic protein (*Mbp*), myelin associated oligodendrocyte basic protein (*Mobp*), brain enriched myelin associated protein 1 (*Bcas1*), myelin associated glycoprotein (*Mag*), proteolipid protein 1 (*Plp1*), and oligodendrocytic myelin paranodal and inner loop protein (*Opalin*); genes related to myelin and oligodendrocyte regulation and development including myelin oligodendrocyte glycoprotein (*Mog*), myelin and lymphocyte protein (*Mal*), myelin regulatory factor (*Myrf*), NK6 homeobox 2 (*Nkx6-2*) and cyclic nucleotide phosphodiesterase (*CNP*); as well as genes involved in myelin-associated signaling and interactions like lysophosphatidic acid receptor 1 (*Lapr1*), tetraspanin-2 (*Tspan2*), contactin 2 (*Cntn2*) and rho guanine nucleotide exchange factor 10 (*Arhgef10*). Further, the expression of many of these genes was relatively higher in reorganized dominants that maintained their status compared to control dominants, indicating that those dominants that maintain status may up-regulate the expression of myelination related genes. Several studies have demonstrated that chronic social stressors can induce reduction in myelin related gene expression followed by longer-term reductions in oligodendrocyte number and myelin protein levels and thickness. Chronic social defeat for up to two weeks is associated with reduction of myelination particularly in the mPFC, hippocampus and nucleus accumbens (Bonnefil et al., 2019; Lehmann et al., 2017), although chronic social stress can also lead to increases in myelination in a brain region specific manner (Poggi et al., 2022). There is also evidence that social stressors lead to reduced expression of oligodendroycte genes encoding myelin and myelin-axon-integrity proteins in the basolateral and central amygdala (Cathomas et al., 2019). Our data extend these findings, suggesting that reductions in myelination processes may already begin to emerge within 70 minutes of a stressful social challenge. Although it is unclear if our observed transcriptional changes lead to long-term structural changes in myelin, our findings suggest that changes in myelination may underlie rapidly learning to avoid socially dominant individuals. Notably, we did not observe differences in myelination gene expression between any group of subordinate animals, suggesting that reductions in myelination may be specific to losing social status.

A profound difference between socially descending animals and those that maintained their dominance status was the lower expression of genes related to cholinergic function in descending animals. 7 out of the top 25 most differentially expressed genes between descenders and dominants were related to this signaling pathway. Animals that maintain their dominance had higher expression of *Chat* which encodes the choline acetyltransferase enzyme, responsible for catalyzing the synthesis of acetylcholine in neurons; the vesicular acetylcholine transporters (*Slc18a3*, *Slc5a7*, *Slc10a4*), integral membrane proteins responsible for packaging and transporting acetylcholine into synaptic vesicles within cholinergic neurons; the muscarinic receptor *Chrm2; Ache* which produces the enzyme acetylcholinesterase that rapidly degrades and inactivates acetylcholine in the synaptic cleft. Further, animals that remained dominant had significantly higher expression of the homeobox genes, *Gbx1* and *Lhx8* that are regulators of cholinergic neuron development (Asbreuk et al., 2002; Mori et al., 2004). *Lhx8* expression also induces Vesicular ACh transporter activity and acetylcholine release (Li et al., 2014; Tomioka et al., 2014). Cholinergic signaling in the amygdala is necessary for social recognition (Prado et al., 2006), social memory formation (Kljakic et al., 2021; Winslow & Camacho, 1995), social stress responsivity (Mineur et al., 2016) and fear learning (Crimmins et al., 2023). Although cholinergic innervation of the medial amygdala is not as extensive as that of other nuclei of the amygdala (Hecker & Mesulam, 1994), our findings strongly suggest that cholinergic activity in the MeA may promote resilience to social challenges in dominant animals.

In contrast to socially descending animals, fewer genes and pathways were found to be uniquely differentially regulated in socially ascending animals compared to reorganized subordinates who remained subordinates. Using WGCNA, we identified a small gene co-expression network (orange module) related to mononuclear cell and lymphocyte differentiation that showed higher expression in maintaining subordinate status compared to ascenders and control subordinates. Previously, social defeat stress has been shown to impact immune regulation in the amygdala and such activation may underlie changes in social memory formation (Munshi et al., 2020; Weber et al., 2017; Wohleb et al., 2011).

### Transitioning social status is associated with shared patterns of gene expression

Within group-living organisms, individuals rarely maintain the same social rank over their entire lifetime. Individuals can ascend and descend in social status due to multiple factors including alternative reproductive tactics (Taborsky & Oliveira, 2012), seasonal changes (Liddle et al., 2022), onset of migratory behavior (Birnie-Gauvin et al., 2021; Gu et al., 2024) or changes in the social or ecological environment. This type of social plasticity can be viewed as major transitions between life history stages, with dramatic changes in behavior, physiology, and gene expression (Gebhardt & Stearns, 1992; West-Eberhard, 2003). It has long been hypothesized that there are dedicated molecular pathways – or “plasticity genes” – that might govern these life history transitions independent of the specific transition in question or its direction. However, evidence in support of this hypothesis has been limited, and the majority of the work has been focused on reproductive efforts and migration patterns. For example, Aubin-Horth et al., (2009) identified in the brains of male Atlantic salmon, a set of candidate genes involving melanocortin, gonadotropin, and growth hormone signaling that are regulated in similar magnitude but opposite direction in the transition from, on the one hand, immature males to sneaker males and, on the other, the difference between early and late migrants to saltwater. Similarly, variation in expression of “plasticity genes” has been argued to underlie the differential susceptibility of individuals to environmental influences (Belsky et al., 2009). The same set of genes may both make some individuals more vulnerable to psychopathology as well as benefiting other individuals when exposed to advantageous environmental contexts such as social support or enrichment.

Our study is the first to characterize transcriptome changes in response to transitions in social status. We found that only 279 genes were differentially expressed in both descending animals compared to dominants <u>and</u> ascending animals compared to subordinates. Significantly, 265 of these genes were differentially expressed in the same direction in both ascenders and descenders, with 142 genes being up and 123 being down in transitioning animals. This suggests that there is a coordinated suite of gene expression changes that is specifically associated with changing social rank. Two genes were extremely overexpressed in transitioning animals (both ascending and descending animals) compared to those that maintained status. These were Myosin binding protein C1 *(Mybpc1)* and μ-Crystallin (*Crym*). Both genes were also identified by our WGCNA as hub genes for the brown module of genes that are significantly elevated in transitioning animals. *Mybpc1* had the largest log fold change in both descenders and ascenders. *Mybpc1* is a molecular marker for a subpopulation of vasoactive intestinal polypeptide-expressing (VIP) interneurons in the cortex (Jiang et al., 2023), but little is known about the function of this neuropeptide in the amygdala. VIP modulates numerous social behaviors including aggression (Kingsbury & Wilson, 2016) as well as stress coping during social stressors like defeat stress (Mineur et al., 2022). Notably, we found that *Vip* was one of the most highly expressed genes in descenders and Vip receptor1 (*Vipr1*) was also overexpressed in ascenders, suggesting that modulation of this sub-population of VIP neurons may regulate social status transitions in either direction.

*Crym* encodes a protein that regulates the amount of freely available thyroid hormone Triiodothyronine (T3) in cytosol (Aksoy et al., 2022). *Crym* has recently been shown to be a critical modulator within the medial amygdala of driving transcriptional changes in response to social isolation stress and the effects of this stress on reward related behaviors (Walker et al., 2022). Importantly, over-expression of *Crym* is associated with increased transcriptional plasticity of the MeA. Consistent with this, we found that both descenders and ascenders had higher expression of the receptor for thyrotropin-releasing hormone receptor (*Trhr*). Moreover, descending males had much higher expression of thyroid releasing hormone as well as thyroid hormone receptors compared to dominant males that did not descend. Previous studies have described a role for thyroid hormone signaling in the amygdala in regulating fear learning (Bárez-López et al., 2017; Montero-Pedrazuela et al., 2011) in the MeA specifically in modulating social vigilance (Kwon et al 2021 Nature). Our findings suggest that altered thyroid hormone signaling, possibly induced via activation of *Crym*, may occur during social status transitions – and maybe life history transitions more generally – to facilitate context-appropriate social behavior.

The hub gene with the highest module membership of the brown module was *Grin2b* (glutamate ionotropic receptor NMDA type subunit 2B), which was significantly more highly expressed in both descenders and ascenders compared to dominants and subordinates. Other genes that regulate the activity of NMDA and AMPA receptors were also overexpressed in transitioning animals, including *Nsmf* (NMDA receptor synaptonuclear signaling and neuronal migration factor) which modulates NMDA receptor activity and downstream signaling pathways, *Gria3* (glutamate ionotropic receptor AMPA type subunit 3) and *Rin1* (Ras and Rab interactor), which controls the activity of AMPA receptors and therefore influences synaptic plasticity associated with fast excitatory neural transmission (Hausser & Schlett, 2019). This upregulation of glutamatergic signaling is unlikely to be related to increased aggression as previous work has demonstrated that activation of GABAergic rather than glutamatergic neurons in the MeA is associated with onset of male aggression (Hong et al., 2014). However, given glutamatergic neurons in the MeA receive projections from olfactory areas (Lu et al., 2023), our findings likely reflect an olfaction-mediated response to learning novel social cues and contexts.

Using a stricter comparison, we found five genes (*Car12, Unc13a, Lpl, Tesc, Pitpnm2*) showing higher expression in both ascenders and descenders compared to both control animals and animals that maintained their ranks. There is evidence that four of these genes play a significant role in regulating synaptic plasticity and learning. Tescalin (*Tesc*) expression leads to dendrite branching and neurite outgrowth (Takamatsu et al., 2017). phosphatidylinositol transfer protein membrane associated 2 (*Piptnm24* plays a role in neurite elongation and synapse formation (Schmidt & Dikic, 2005). *Unc13a* expression in mushroom bodies plays a fundamental role in olfactory memory encoding in Drosophila (Pooryasin et al., 2021), and hippocampal expression of lipoprotein lipase (*Lpl*) is necessary for synaptic vesicle recycling and acquisition of memories in mice (Kim et al., 2014).

## Conclusion

In the present study, we developed a novel social reorganization paradigm to investigate behavioral, physiological, and molecular adaptations in male mice that socially ascend or descend within dominance hierarchies. We demonstrate that, following a social disruption, outbred CD-1 mice rapidly reorganize themselves into linear dominance hierarchies, even in animals with a history of being socially subordinate. Previously dominant animals showed the largest response to this social reorganization, exhibiting significantly higher levels of plasma corticosterone and a two-fold increase in DEGs in the MeA, a central hub of the SDMN. Within the MeA, animals that maintained their high dominance status had relatively higher expression of genes related to cholinergic signaling and myelination compared to descending males. Additionally, we identified novel social transition genes that show changes in expression in both socially descending and ascending animals. Two of these genes, *Mybpc1* and *Crym* are exceptionally highly expressed in animals that socially ascend or descend suggesting that they coordinate the activity of several other genes such as those that regulate thyroid hormone signaling. WGCNA also revealed that genes associated with synaptic plasticity and glutamatergic signaling are highly expressed in status transitioning animals, which may be relevant for learning of novel social relationships. Overall, our results demonstrate the existence of some common molecular changes that are associated with rapid social ascent and descent in the MeA, suggesting that this critical node of the SDMN is remodeled as animals learn to navigate new social environments.

## Materials and Methods

### Animals, Housing and Behavioral Observations

Male CD-1 mice (N = 260) aged 7-8 weeks were obtained from Charles River Laboratory (Houston, TX, USA) and housed at The University of Texas at Austin. The animal facility was kept on a 12/12 light/dark cycle, with white light at 2300 h and red lights (dark cycle) on at 1100 h with constant temperature (21–24 °C) and humidity (30–50%). Upon arrival, animals were randomly placed into 65 groups of four animals for 10 days in two standard rat cages (35.5 x 20.5 x 14cm) covered in pine shaving bedding and connected by Plexiglass tubes with multiple enrichment objects (**Supplemental Figure 1).** Standard chow and water were provided ad libitum. Mice were marked with a nontoxic animal marker (Stoelting Co., Wood Dale, IL, USA) to display unique identification on group housing day 3 or 4 (GD3/4). Live behavioral observations began on GD3/4 and were conducted daily until GD10 or GD11 during the dark cycle by a trained observer, as described previously (So et al., 2015; Williamson et al., 2016).Briefly, all occurrences of agonistic (fighting, chasing, mounting) and subordinate behaviors (freezing, fleeing, subordinate posture) between two individuals were recorded using either an Android device or Apple iPad and directly uploaded to a timestamped Google Drive via a survey link. On average, one hour per day of live observation were conducted per group, resulting in an average of 82 agonistic/subordinate behavioral interactions per group. All procedures were conducted with the approval of the Institutional Animal Care and Use Committee of the University of Texas at Austin.

### Submandibular Blood Collection

At Zeitgeber time 01 (ZT01) on GD7/8, 100 to 200 µl of blood was collected into EDTA-coated Vacutainer tubes (Becton Dickinson) from each mouse via submandibular puncture as described in (Golde et al., (2005). Once blood collection was completed 2-3s after the punch, sterile gauze was immediately used to stop bleeding. Blood samples were gently inverted a few times, stored on ice for 45–60 min, and then centrifuged (1500xg for 15 min at 4°C). Plasma was collected and held at -80°C until the corticosterone assay was completed.

### Body Mass Assessment

Individual body mass was monitored throughout the study. Mouse mass was collected on GD3/4 as mice were individually marked. Mice were weighed again, before and after blood collection, on GD7/8. Final body mass was measured right before tissue collection on GD10 or GD11 after the social reorganization paradigm was completed.

### Experimental Reorganization of Social Hierarchy

At the start of GD10, between ZT23.5 and ZT00, mice from stable social hierarchies were randomly assigned to reorganized (N=52 groups) or control groups (N=13 groups). Mice undergoing social reorganization were placed into new groups of four with unfamiliar animals of equal social status as determined by their rank in their pre-reorganization group. Therefore, there were four reorganized conditions: groups consisting of i) four unfamiliar alpha (rank 1) males, ii) four unfamiliar beta (rank 2) males, iii) four unfamiliar gamma (rank 3) males, or, iv) four unfamiliar delta (rank 4) males (**Figure 1A**). Animals remained in reorganized groups for either 70 min (N=28 groups) or 25 hours (N=24 groups). In the control condition, all four animals in each group were placed into a new cage with fresh bedding. Mice were housed for 70 min (N= 6 groups) following the cage change and another set of mice were kept for 25hrs (N= 7 groups) following the cage change. Live behavioral observations were conducted for all groups during the 70 min immediately following social reorganization (or cage change for the control condition) using the same procedure as used for stable hierarchies. Additionally, for animals that were housed for 25hrs, 3 hours of behavioral observations were conducted directly following cage changes, with an additional 1 to 2 hours during the 25hrs period.

### Tissue Collection and Preparation

70 min after social reorganization, mice were euthanized via rapid decapitation. Trunk blood samples were collected into EDTA-coated Vacutainer tubes (Becton Dickinson) and gently inverted a few times and stored on ice for 45–60 min before being centrifuged (1500xg for 15 min at 4°C) for plasma collection and stored at −80 °C until the corticosterone assay was completed. Brains were collected and flash-frozen in a hexane cooling bath, placed on dry ice and stored at -80 until further processing. Brains from animals who socially ascended (ASC, rank 4 to rank 1, N_70min_ = 7), descended (DES, rank 1 to rank 4 N_70min_ = 7), remained dominant (DOM, rank 1 to rank 1 N_70min_ = 7), and remained subordinate (SUB, rank 4 to rank 4, N_70min_ = 7) during reorganization as well as control dominant (CDOM, rank 1 to rank 1 N_70min_ = 6).

Control subordinates (CSUB, rank 4 to rank 4, N_70min_ = 6) were selected for Tag-based RNA sequencing (Lohman et al., 2016; Meyer et al., 2011a). These whole brains were sectioned on a cryostat (Leica Biosystems, Deer Park, IL) at 300µm, then the Medial Amygdala (MeA) was dissected separately using a Stoelting 0.51 mm tissue punch (Stoelting, Wood Dale, IL, Cat. No. 57401) and homogenized in 100ul lysis buffer (Thermo Fisher Scientific, Waltham, MA; MagMax Total RNA isolation kit, Cat. No. A27828) with 0.7% beta-mercaptoethanol. Lysates were further processed on KingFisher Flex (Thermo Fisher Scientific, Cat. No. 5400630l) for total RNA isolation. RNA quality was determined using RNA 6000 Nano Assay with BioAnalyzer, processed samples had an average RIN of 8.5, well above the 7 cutoff (Agilent Technologies, Santa Clara, CA). RNA concentration was determined with Quant-it RNA High Sensitivity assay kit (Thermo Fisher Scientific, Cat. No. Q33140). RNA samples were normalized to 10 ng/µl and stored at -80°C before sequencing. In total, 40 extracted RNA samples (N = 7/reorganized groups (ASC, DES, DOM, SUB), N= 6/control groups (CDOM, CSUB) were submitted and processed in the same batch at the Genome Sequence and Analysis Facility at the University of Texas at Austin for Tag-based RNA sequencing. Libraries were constructed according to (Lohman et al., 2016). Reads were sequenced on the NovaSeq 6000 SR100 with minimum reads of 4 million, and the target reads per sample of 5 million.

### Corticosterone Enzyme Immunoassay

Plasma obtained, on GD7, prior to and both at 70 min and 25 hr after the social reorganization was selected from mice who socially ascended (ASC, rank 3 to rank 1, N_70min_ = 14, N_25hr_ = 11; rank 4 to rank 1, N_70min_ = 14, N_25hr_ = 12), descended (DES, rank 1 to rank 4 N7_0min_ = 14, N_25hr_ = 11; rank 1 to rank 3 N_70min_ = 14, N_25hr_ = 12), remained dominant (DOM, rank 1 to rank 1 N_70min_ = 14, N_25hr_ = 12), and remained subordinate (SUB, rank 3 to rank 3, N_70min_ = 14, N_25hr_ = 11; rank 4 to rank 4, N_70min_ = 14, N_25hr_ = 10). Plasma was also obtained from control dominant (CDOM, rank 1: N_70min_ = 12, N_25hr_ = 12) and subordinate (CSUB, rank 3: N_70min_ = 12, N_25hr_ = 13; rank 4: N_70min_ = 12, N_25hr_ = 12) mice. Plasma corticosterone concentrations were measured using a commercially available ELISA kit (Arbor Assays, Ann Arbor, MI). The ELISA was carried out by the manufacturer’s instructions using a 96-well plate reader with absorbance at 450nm.

## Statistics

All statistical analyses were carried out in R version 4.2.1 (R Core Team, 2022).

### i. Behavioral and Hormonal Data

Win/loss sociomatrices for each social group were created both pre- and post-reorganization using the frequency of wins and losses observed from behavioral observations (**Supplemental Figures 2 & 3).** From these sociomatrices, we tested the stability of each social hierarchy by testing the significance of directional consistency as previously described (Williamson et al., 2016, 2019). Individual social ranks were determined by David’s Scores (DS). DS are a measure of individual dominance related to the proportion of wins to losses adjusted for opponents’ relative dominance (Gammell et al., 2003). DS were calculated pre-reorganization and at 70 min and 25hrs after reorganization using the R package ‘compete’ (Curley, 2016). To test for differences in the mean DS for each rank pre-reorganization between groups that were assigned to the reorganized and control conditions we used a One-Way ANOVA with group-condition as the predictor. We also tested for differences in the mean DS for each rank post-reorganization between groups consisting of all previously alpha, beta, gamma or delta males using previous status as the predictor. This was done using One-Way ANOVA with Tukey post-hoc comparisons. For all One-Way ANOVAs data were normal and homogeneity of variance was determined. We used generalized linear mixed-effect models (GLMMs) to determine what predicted DS before or after reorganization. We conducted a GLMM with pre-reorganization DS as the outcome variable, and body mass at GD3/4 or GD7/8 as a fixed effect, along with individual id, batch, and group id as random effects to test if body mass was associated with dominance prior to reorganization. Post-reorganization, we conducted separate GLMMs for each group type (i.e. those consisting of all previously alphas, betas, gammas or deltas). Each GLMM had post-reorganization DS as the outcome variable and the fixed effect being each individual’s body mass at GD10, pre-reorganization DS, rate of aggression given pre-reorganization, or rate of aggression received pre-reorganization. Rates of aggression given and received were calculated as the total number of aggressive acts given or received per hour by each individual. We also included a fixed effect ‘total aggression rate per group’, which was the rate per hour of all aggression in each individual’s group prior to reorganization. This was to test if behavioral dynamics within the group-of-origin influenced post-reorganization DS. Random effects were individual id, batch and group id. To compare if groups of all alphas, betas, gammas and deltas differ in their behavior post-reorganization we ran GLMMs with either latency to fight or total number of fights in the 70 minutes post-reorganization as outcome variables. The fixed effect was the group-status (i.e., whether animals were previously all alphas, betas, gammas or deltas). Random effects were batch and group id.

We performed a randomization test to determine if individuals that were housed together pre-reorganization were more likely than chance to attain similar ranks in their post-reorganization hierarchies. For each of the four animals within each group, we determined the sum of the differences in their post-reorganization ranks. For example, if all animals achieved the same rank, then the sum of rank differences would be 0. However, if two animals achieved rank 1, and two animals achieved rank 4, then the sum of the pairwise rank differences is 12. We calculated the total sum of all pairwise rank differences across all groups and compared this observed value to the theoretical distribution of possible pairwise rank differences across groups assuming that ranks achieved are independent of prior group membership. We performed this randomization 10,000 times and calculated the p-value by determining the proportion of randomizations that showed a sum of pairwise rank differences across all groups higher than our observed value.

Using Binomial tests, we examined whether post-reorganization hierarchies were stable between the first 70 minutes of observation and 25 hours by testing whether the proportion of animals of each rank (as calculated after the first 70 minutes) that went on to win or lose more fights against animals of relatively higher or lower ranks was less than expected by chance. We used a GLMM with a Gamma likelihood and log link including plasma corticosterone (CORT) as the outcome variable, social status or group condition (control vs. reorganized) as fixed effects, and id, group id, batch, and plate as random effects, to test the relationship between pre-plasma CORT levels and social status or experimental group membership prior to reorganization. To examine the relationship between the change in CORT between pre-reorganization and post-reorganization, we ran a linear mixed effect model (LMM) with the difference in CORT between the two time points as the outcome variable, social condition (remaining dominant, descending, ascending, remaining subordinate or control) as a fixed effect, with group-ids pre- and post-reorganization as random effects. Appropriate GLMMs and LMMs were used for each analysis according to the data distribution and residuals from fitted models using the R package ‘lme4’ (Bates et al., 2023). We used the package ‘lmeRTest’ (Kuznetsova et al., 2020) to derive p-values for GLMMs and LMMs.

### ii. Tag-based RNA Sequencing Analysis

Raw reads were processed and mapped to the *Mus musculus* reference genome (Ensembl release 99) using bowtie with an average mapping rate of 73.6 % (Langmead & Salzberg, 2012) to obtain gene count data by following the Tag-Seq data processing pipeline (Lohman et al., 2016; Meyer et al., 2011), as described in (Lee et al., 2022). Briefly, customized perl script utilizing FASTX-Toolkits (Hannon, 2010) CUTADAPT v. 2.8 (Martin, 2011) was used to remove reads with a homo-polymer run of “A”≥8 bases and retain reads with minimum 20 bases and removal PCR duplicates. Quality of raw sequence data was checked with FastQC (Andrews, 2010). All samples passed quality control and were used in the analysis.

*Analysis of differentially expressed genes (DEGs):* All DEG analyses were conducted using the Bioconductor package ‘*limma’* (Smyth et al., 2021). We first filtered out genes with <10 counts across all 40 samples. Next, filtered read counts were normalized by the trimmed mean of the M-values normalization method (TMM, (Smyth et al., 2021). Two different DEG analyses were conducted with filtered normalized count data with a voom transform. First, our analysis identified DEGs in previously dominant and control dominant animals with 3 comparisons: descending (DES) vs. dominant (DOM) males, DES vs. control dominant (CDOM), and DOM vs. CDOM males. Second, we identified DEGs in previously subordinate and control subordinate individuals with 3 comparisons: ascending (ASC) vs. subordinate males (SUB), ASC vs. control subordinates (CSUB), and SUB vs. CSUB. Raw p-values for each DEG analysis were adjusted via empirical false discovery rate (eFDR, (Storey & Tibshirani, 2003).

We permuted sample labels 5000 times and obtained a null distribution of p-values to estimate the empirical false discovery rate. We set a threshold for differentially expressed genes as 20% change of the absolute values of log2 fold change at the empirical false discovery rate (eFDR) of 5%. Additionally, we hypothesize that animals that transitioned social status would show relatively higher expression of IEGs compared to those that did not transition. To test this, we compared whether a significantly higher proportion of these genes that are DEGs were relatively more highly expressed in DES vs DOM and ASC vs SUB comparisons using Binomial Tests.

We adapted a curated list of genes from (Tyssowski et al., 2018) to only include rapid and delayed response genes. To examine common DEGs between DES vs DOM and ASC vs SUB comparisons, we created a 2x2 matrix with cells corresponding to the total number of DEGs that are more highly or lowly expressed in each group. We used a chi-squared test to determine if cells had distributions significantly different from random, and calculated the phi-coefficient to identify if DEGs were directionally consistent across groups in their expression. Enriched gene ontology (GO) analysis was conducted to explore functional differences among the different comparisons of DEGs. We used the *‘clusterProfile*’ R package (Yu et al., 2020) to test the statistical overlap between annotated biological and functional significance in each DEG set.

*Weighted gene co-expression analysis (WGCNA)* WGCNA was performed using the *‘WGCNA’* R package (Langfelder & Horvath, 2008). For the first WGCNA analysis, we combined count data across all social condition (reorganized and control animals of both dominant and subordinate status) to construct one gene co-expression network. Combining all social conditions led to 40 samples, above the recommended sample size to use with this analysis. WGCNA modules were constructed in signed hybrid mode with a minimum module size set at 50 genes. To investigate social transitions, we conducted a second WGCNA excluding control animals. We pooled ascending and descending males into one group of transitioning (TRN) animals to compare to reorganized dominant and subordinate males that maintained their status. WGCNA modules were constructed in signed hybrid mode with a minimum module size set at 100 genes. To construct each of these signed hybrid correlation networks power values were set at 4, based on the scale-free topology criterion (**Supplemental Figure 13** , Zhang & Horvath, 2005). Gene counts were normalized with the limma (Smyth et al., 2021) package, and genes that had low expression counts (<10) across all samples were filtered out prior to network construction (**Supplemental Figure 13**). WGCNA co-expression modules were associated with social condition via Pearson correlation matrices, which clustered highly co-expressed genes into several modules (**Supplemental Figure 13**). After, identifying modules of co-expressed genes within the network, we calculated the module eigengene (ME). ME is the first principal component of the gene expression levels of all genes contained in the module which summarizes overall gene expression profile. Further in modules of interest, we calculated module membership (MM) for each gene. MM is measured as the Pearson correlation between the gene expression level and the module eigengene. A module membership close to 1 indicates that the gene is highly connected to other genes in the module. Genes with MM above 0.8 were considered hub genes in their modules. A linear mixed effects model was performed with the social condition as the fixed factor and group id as a random factor to determine the association between each module eigengene and social condition. Modules were further characterized into descending, ascending, or social transition modules depending on their differentially expressed module eigengenes (deMEGs) from the LMM. Enriched gene ontology (GO) analysis for genes contained in each module was conducted via the *‘clusterProfiler’* R package (Yu et al., 2020).

## Data Availability Statement

Data and code used in this paper are available at the following repository: https://github.com/ty14/SocialTransitions

