## Supplemental Figures and Tables for "Rapid changes in plasma corticosterone and medial amygdala transcriptome profiles during social status change reveal molecular pathways associated with a major life history transition in mouse dominance hierarchies"

**Supplemental Figure 1:** A) Experimental design. Each procedure was delayed by a day for half the cohorts that went through a social reorganization at 70 minutes to ensure the same start time for all social reorganizations. GD = Group House Day B) Mice are housed in a 4 per cage system (cage lids not shown).

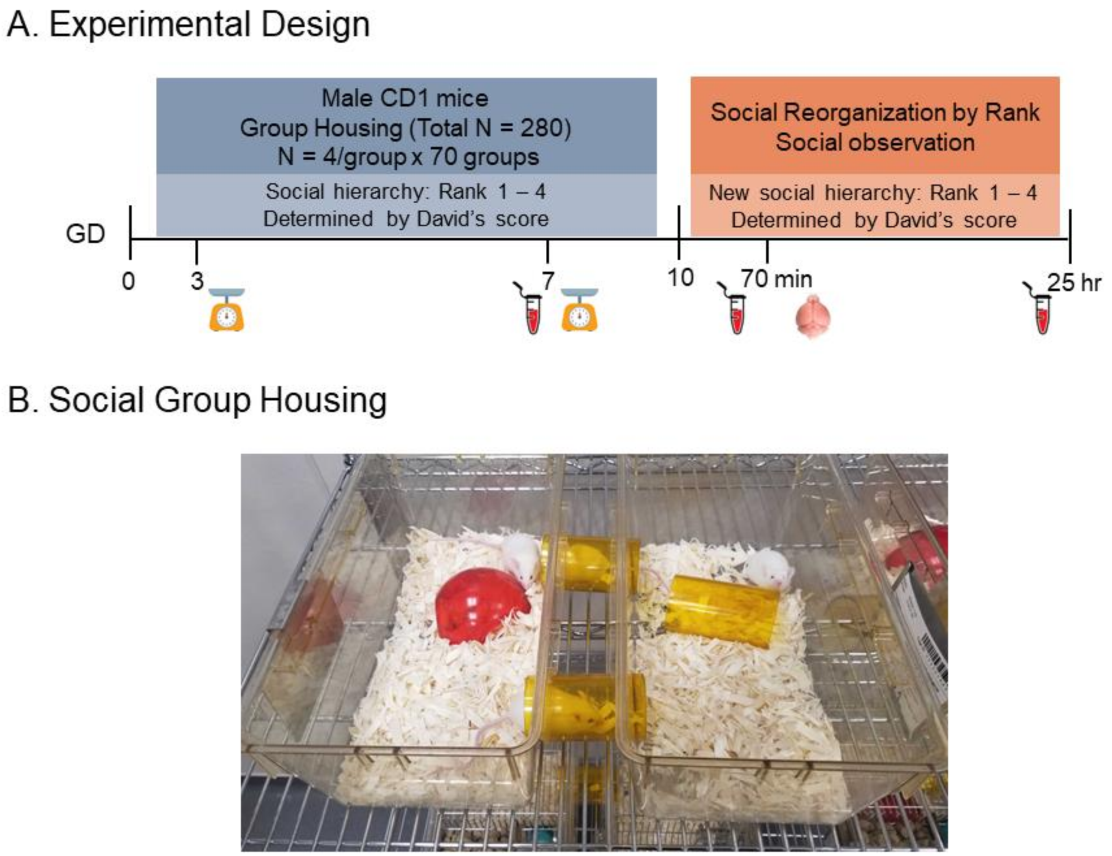

**Supplemental Figure 2:** A) Raw Pre-Reorganized sociomatrices of wins and losses for each cohort. Each value represents the total number of wins by the individual in each row against the individual in each column. The degree of redness represents the frequency of wins. B) Binary matrices for each cohort. A 1 in a cell represents that the individual in the row is a consistent winner against the individual in the column. Numbers on rows and columns refer to animal IDs.

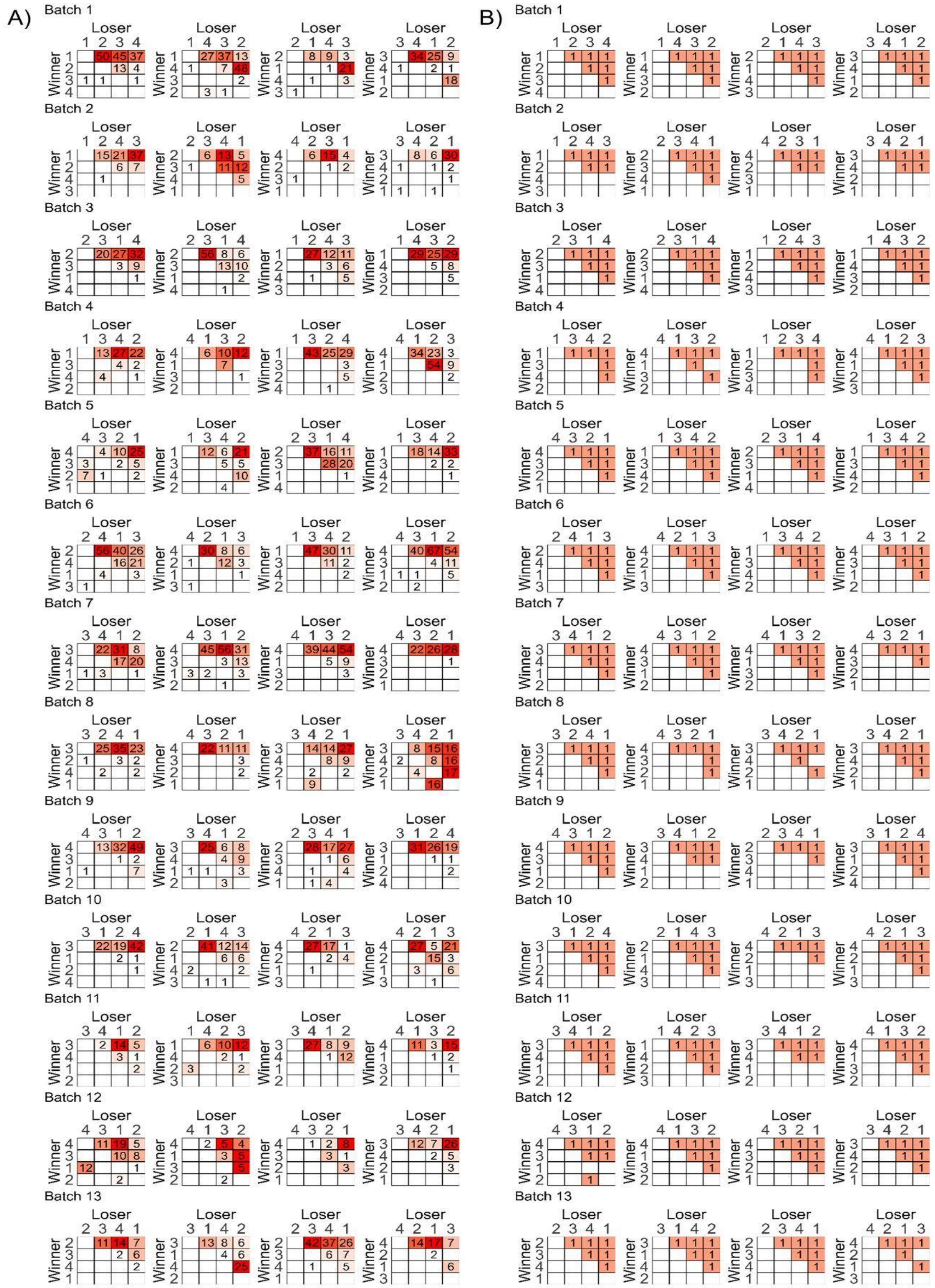

**Supplemental Figure 2:** C) Raw Post-Reorganized sociomatrices of wins and losses for each cohort. Each value represents the total number of wins by the individual in each row against the individual in each column. The degree of redness represents the frequency of wins. D) Binary matrices for each cohort. A 1 in a cell represents that the individual in the row is a consistent winner against the individual in the column. Numbers on rows and columns refer to animal IDs.

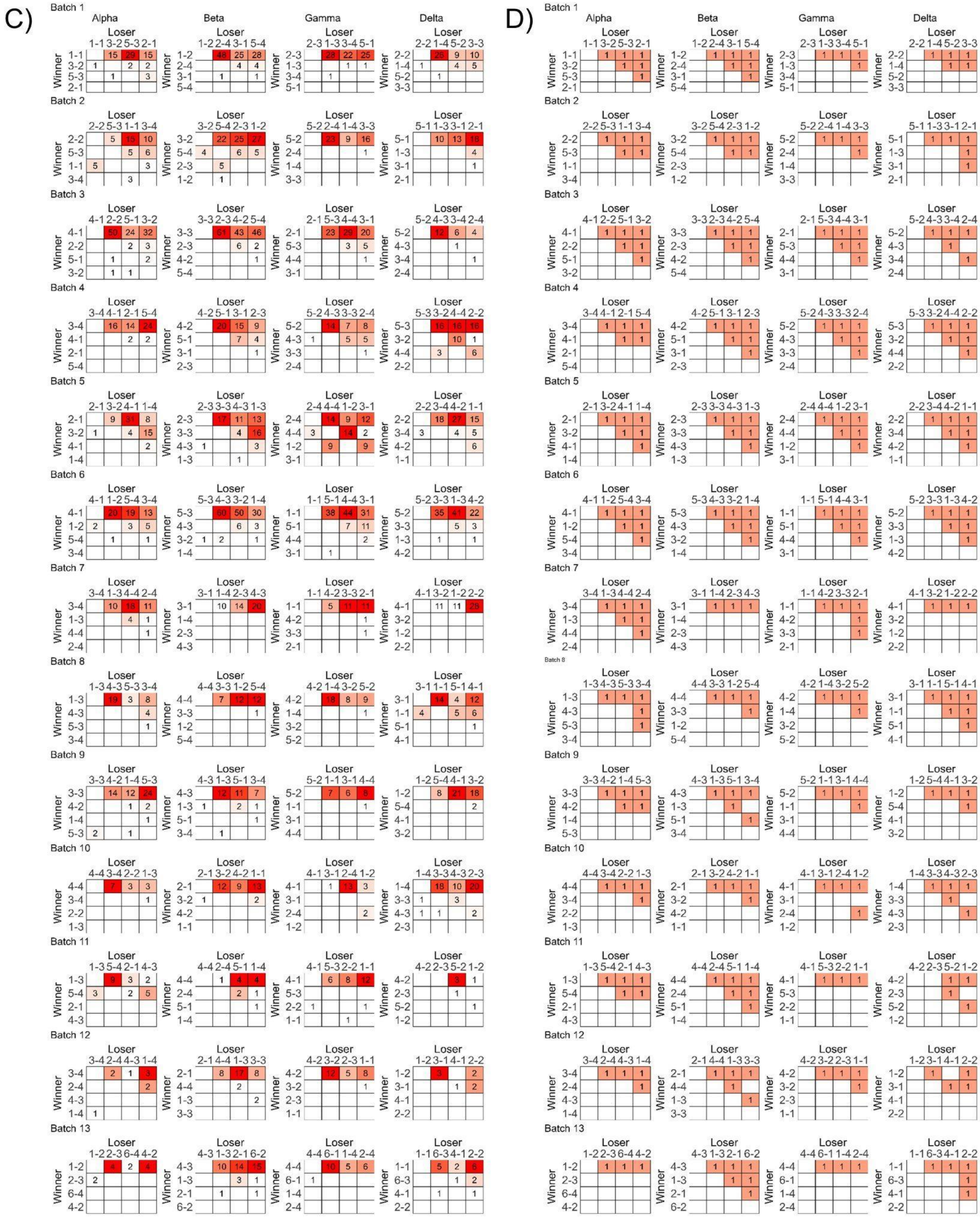

**Supplemental Figure 3:** A) Raw sociomatrices of wins and losses for each cohort of control animals. Each value represents the total number of wins by the individual in each row against the individual in each column. The degree of redness represents the frequency of wins. B) Binary matrices for each cohort. A 1 in a cell represents that the individual in the row is a consistent winner against the individual in the column. Numbers on rows and columns refer to animal IDs.

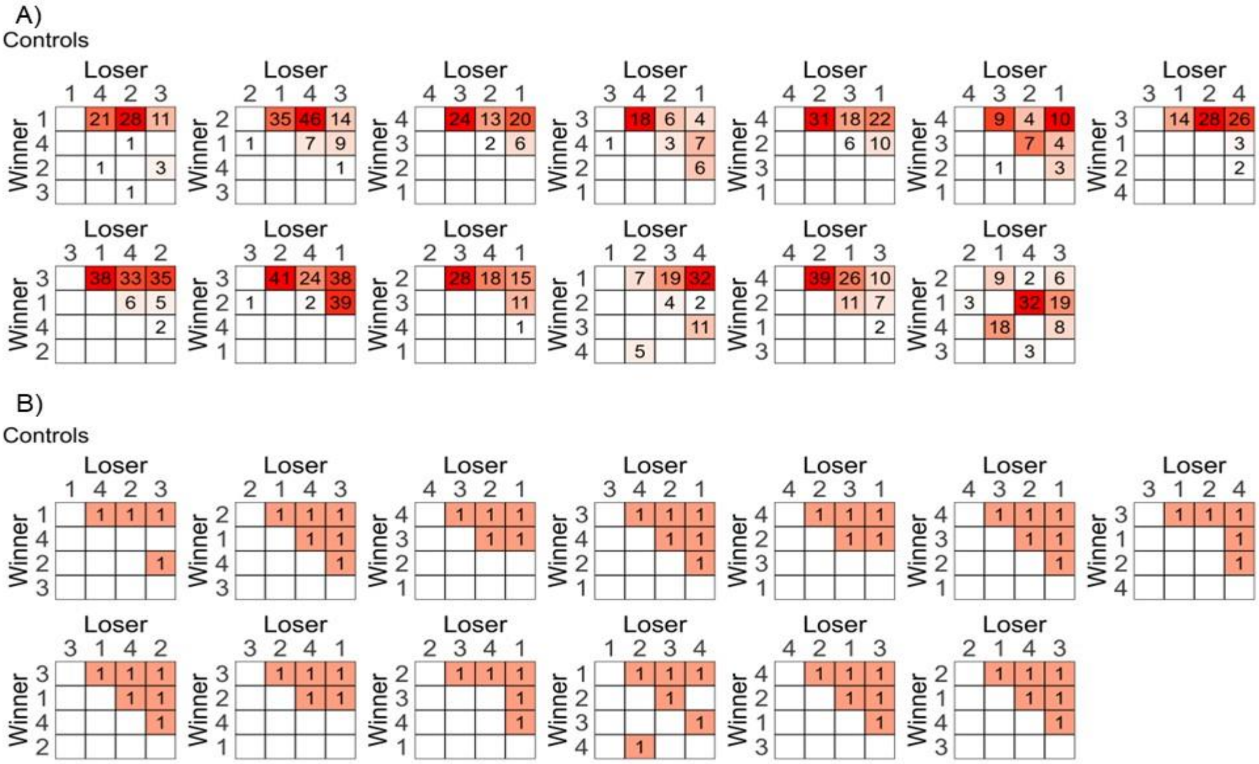

**Supplemental Figure 4:** Boxplots showing median and IQR of body mass by pre-reorganization social rank at the beginning of social observation on day 3 or 4 of group housing (A) and once groups were stable on day 7 or 8 (B) of group housing. Points represent individuals.

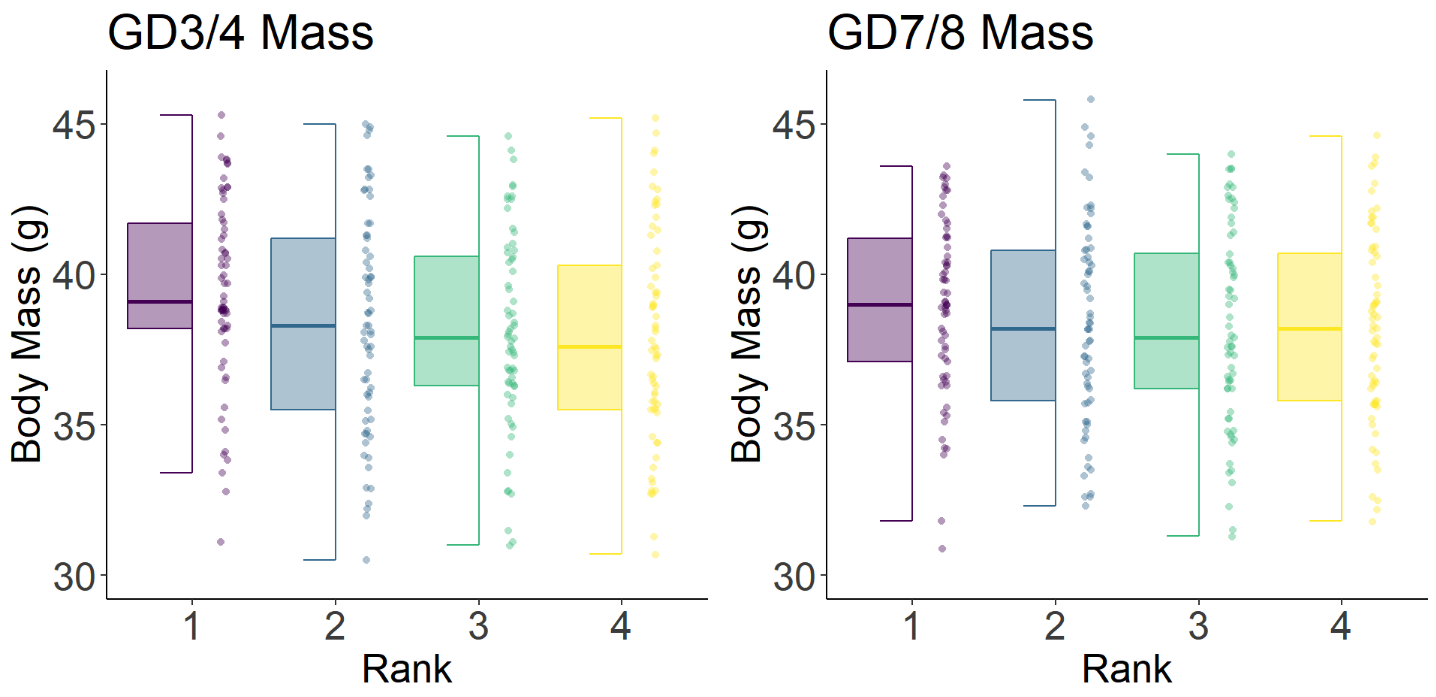


**Supplemental Figure 5**: Total frequency of fights in each group 70 minutes after social reorganization. Boxplots represent medians and IQRs. Points represent individuals.

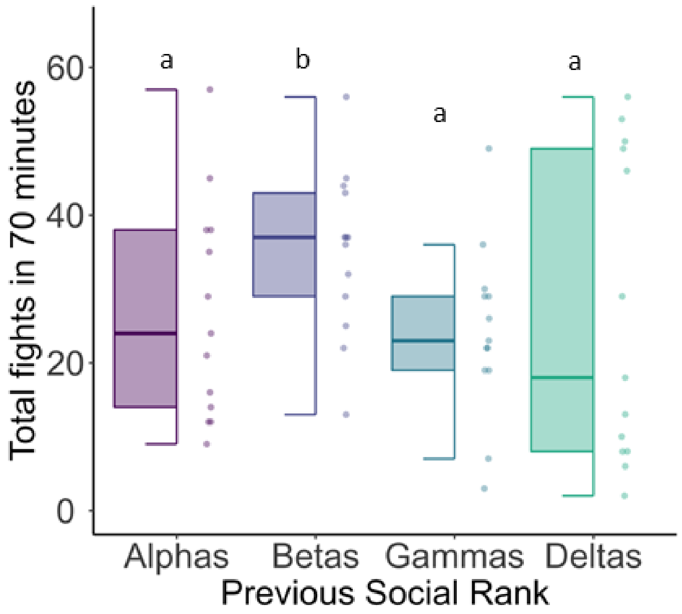

**Supplemental Figure 6**: Correlation of Rates of A) aggression given or received prior to social reorganization (x-axis), or (B) total aggression in each group (x-axis) and post-reorganization David's score (y-axis). Each facet represents individuals in post-reorganization groups of 1) previously alpha males, 2) previously beta males, 3) previously gamma males, 4) previously delta males. Higher rates of aggression in pre-reorganization groups are associated with higher dominance scores post-reorganization in previously beta males. C) Group of Origin leads to similar post ranks. The green distribution is the theoretical distribution of differences in post-reorganization ranks between animals from the same pre-reorganization groups. The dashed line represents the observed differences in post-reorganization ranks.

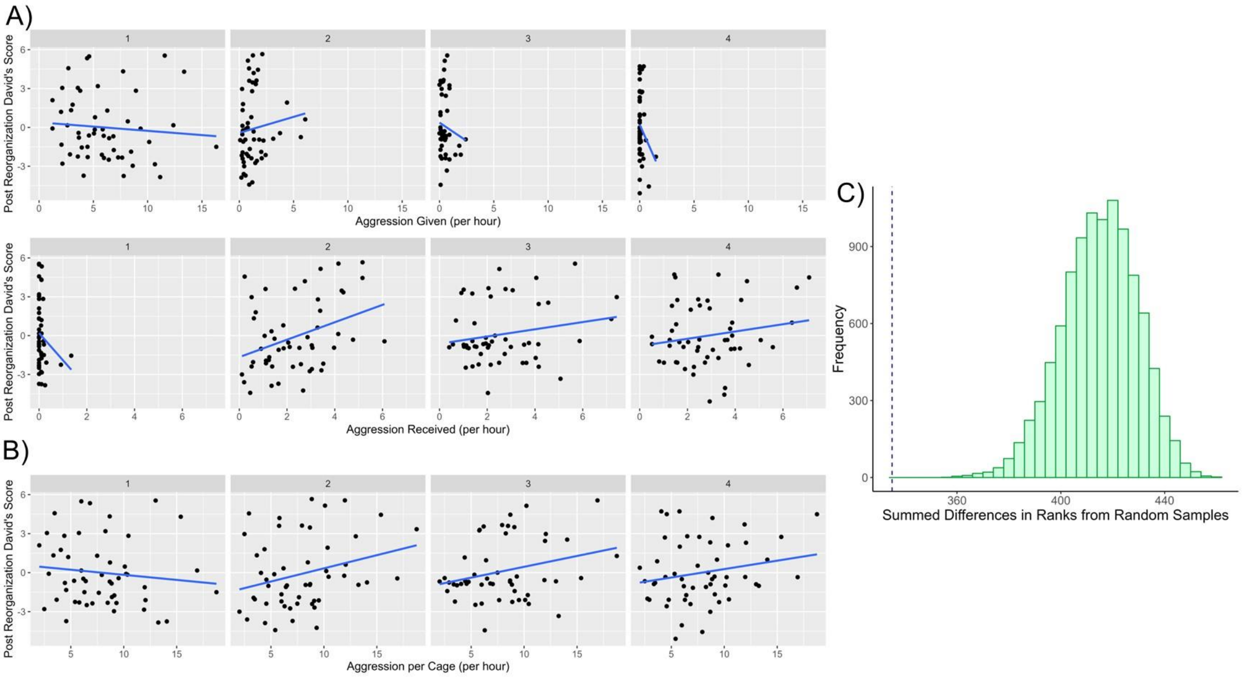

**Supplemental Figure 7:** No significant differences between (A) dominant and subordinate males or (B) control and reorganized males in plasma corticosterone prior to social reorganization. Boxplots represent medians and IQRs. Points represent individual subjects.

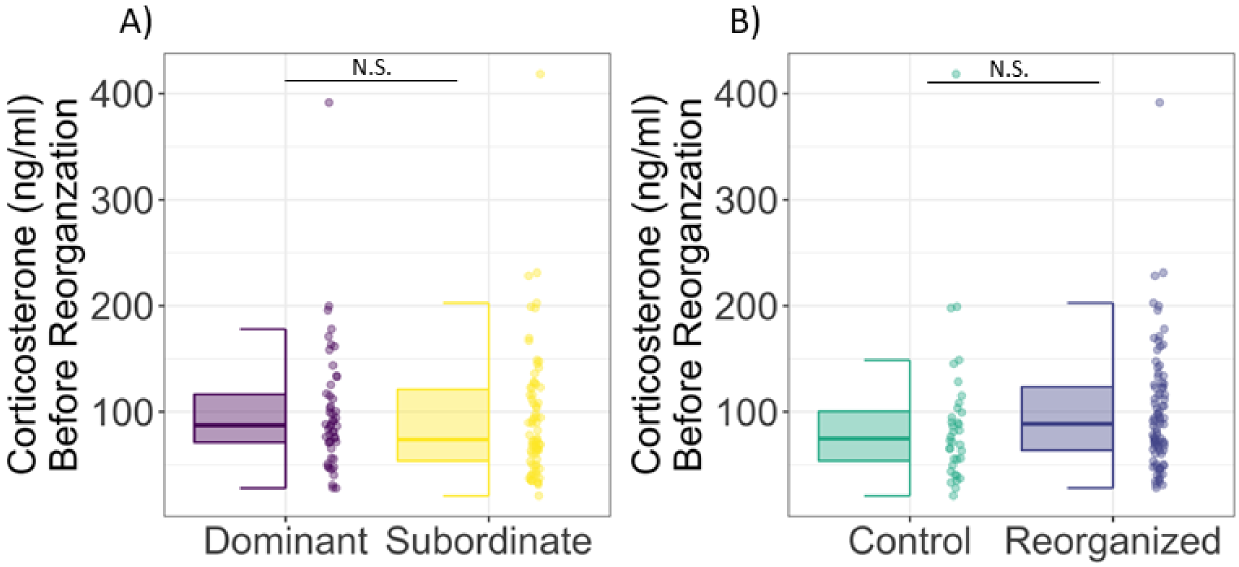

**Supplemental Figure 8**: Volcano plots represent log2 fold change by -log10 of eFDR, 0.05, for each of the comparisons A) DES vs. DOM, B) DES vs. CDOM, and C) DOM vs. CDOM. DOM = previously dominant males that remain dominant; DES = previously dominant males that socially descend; CDOM = control dominant animals that remain dominant. Each volcano plot is annotated with genes with largest fold changes as well as lines indicating log2 fold change at 0.5 and 0.75. D) The total number of DEGs, top ten genes with the largest fold change, and corresponding top biological processes GO-terms**.**

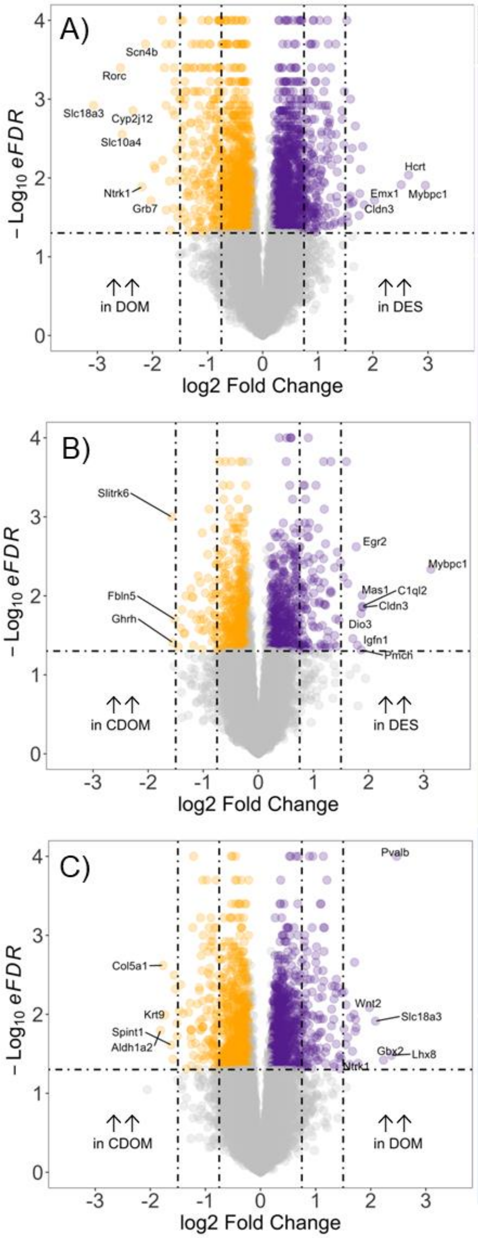


| D) MeA 70min | # of DEGs | Top genes | GO terms |
|---|---|---|---|
| Relatively ↑↑ DES vs. DOM | 1187 | *Mybpc1 Hcrt Emx1 Cldn3 Mas1* *Crym Vip Spink8 Trh Col5a1* | - hormone secretion<br>- regulation of dendrite morphogenesis<br>- peptidyl-serine phosphorylation<br>- regulation of neuron apoptotic process<br>- regulation of synaptic plasticity |
| Relatively ↓↓ DES vs. DOM | 1130 | *Lhx8 Slc18a3 Rorc Slc10a4 Cyp2j12* *Ntrk1 Scn4b Grb7 Skor1 Lrrc38* | - axon ensheathment<br>- myelination<br>- glial cell differentiation<br>- oligodendrocyte differentiation<br>- cellular amino acid metabolic process |
| Relatively ↑↑ DES vs. CDOM | 575 | *Mybpc1 Cldn3 C1ql2 Mas1 Pmch* *Dio3 Igfn1 Egr2 Hcrt Ptgs2* | - regulation of fat cell differentiation<br>- negative regulation of phosphorus<br>- metabolic process<br>- negative regulation of MAPK cascade<br>- cellular response to peptide hormone stimulus |
| Relatively ↓↓ DES vs. CDOM | 526 | *Pvalb Lhx8 Gbx2 Skc18a3 Wnt2* *Ntrk1 Smyd1 Irx2 Chrna6 Scn4b* | - regulation of DNA replication<br>- positive regulation of amino acid transport<br>- regulation of neurogenesis<br>- gliogenesis<br>- myelination |
| Relatively ↑↑ DOM vs. CDOM | 803 | *Slitrk6 Ghrh Fbln5 Obp2b Cbr2* *Sertm2 Aspn Dmrtb1 Tnc Galr1* | - ATP metabolic process<br>- generation of precursor metabolites and energy<br>- energy derivation by oxidation of organic compounds<br>- oxidative phosphorylation<br>- nucleotide metabolic process |
| Relatively ↓↓ DOM vs. CDOM | 832 | *Aldh1a2 Col5a1 Krt9 Spint1 Ccdc42* *Zbrb7c Esyt3 Barx2 Ppp1r17 Hnf1b* | - postsynapse organization<br>- histone modification<br>- positive regulation of DNA repair<br>- neural precursor cell proliferation<br>- I-kappaB kinase/NF-kappaB signaling |

**Supplemental Figure 9:** Volcano plots represent log2 fold change by -log10 of eFDR, 0.05, for each of the comparisons A) ASC vs. SUB, B) ASC vs. CSUB, and C) SUB vs. CSUB. SUB = previously subordinate males that remain subordinate; ASC = previously subordinate males that socially ascend; CSUB = control subordinate animals that remain subordinate. Each volcano plot is annotated with genes with largest fold changes as well as lines indicating log2 fold change at 0.5 and 0.75. D) The total number of DEGs, top ten genes with the largest fold change, and corresponding top biological processes GO-terms.

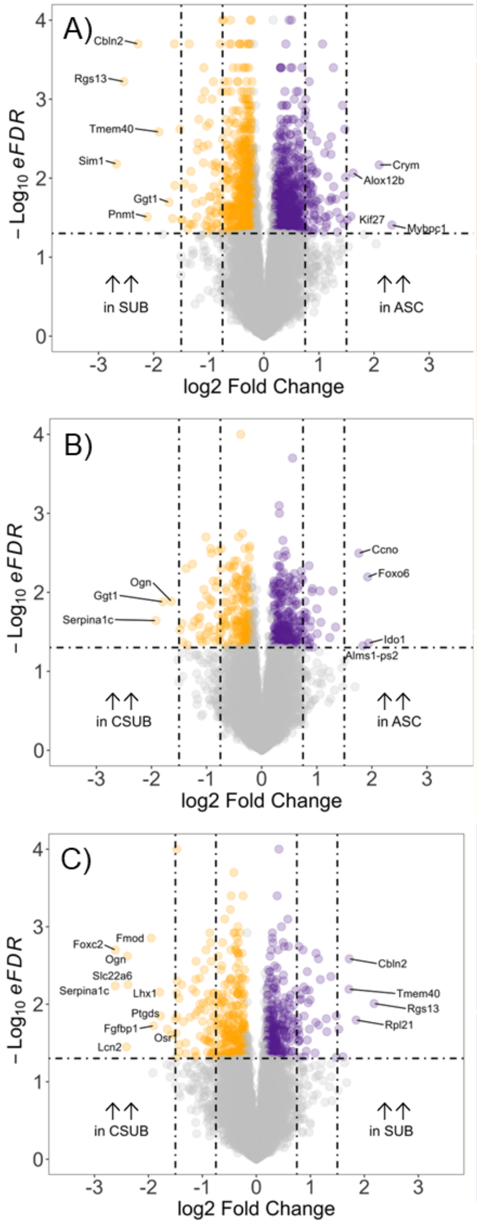


| D) MeA 70 min | # of DEGs | Top genes | GO terms |
|---|---|---|---|
| Relatively ↑↑ ASC vs. SUB | 658 | *Mybpc1 Gpr6<br>Crym1 Ddit4l<br>Alox12b Olfr1393<br>Kif27 C1qtnf2<br>Osr1 Atp6v1c2* | - growth hormone receptor signaling pathway<br>- regulation of body fluid levels<br>- membrane lipid catabolic process<br>- leukocyte migration<br>- positive regulation of lymphocyte differentiation |
| Relatively ↓↓ ASC vs. SUB | 737 | *Sim1 Ggt1<br>Rgs13 Cyp26b1<br>Cbln2 Qrfpr<br>Pnmt Slc6a3<br>Tmem40 C1ql1* | - regulation of neurotransmitter levels<br>- ribonucleoprotein complex biogenesis<br>- regulation of heart contraction<br>- catecholamine metabolic process<br>- locomotory behavior |
| Relatively ↑↑ ASC vs. CSUB | 280 | *Ido1 Tnfrsf25<br>Foxo6 Ano2<br>Alms1-ps2 Gm5127<br>Ccno Mcm5<br>Ttg2 Car12* | - regulation of ion transmembrane transport |
| Relatively ↓↓ ASC vs. CSUB | 225 | *Serpina1c Ptgds<br>Ggt1 Itih2<br>Ogn Mup4<br>Foxc2 Crygb<br>Parpbp Cytl1* | - glycoprotein metabolic process<br>- glycoprotein biosynthetic process<br>- alpha-amino acid metabolic process<br>- protein N-linked glycosylation<br>- cysteine metabolic process |
| Relatively ↑↑ SUB vs. CSUB | 309 | *Rgs13 Gm19410<br>Ppl21 Dkkl1<br>Cbln2 Foxo6<br>Tmem40 Dach2<br>Sim1 Neurod6* | - release of cytochrome c from mitochondria<br>- apoptotic mitochondrial changes<br>- positive regulation of neuron death<br>- peptidyl-lysine modification<br>- regulation of GTPase activity |
| Relatively ↓↓ SUB vs. CSUB | 322 | *Serpina1c Fmod<br>Foxc2 Fgfbp1<br>Lcn2 Lhx1<br>Ogn Ptgds<br>Slc22a6 Osr1* | - positive regulation of response to external stimulus<br>- negative regulation of autophagy<br>- regulation of cytokine-mediated signaling pathway<br>- glycoprotein metabolic process<br>- glycosylation |

**Supplemental Figure 10:** Volcano plots showing log2 fold change and significance (eFDR) for genes in the primary response gene set curated from Tyssowski et al., 2018. DEGs met criteria if the absolute values of log2 fold change were greater than 20% at the empirical false discovery rate (eFDR) of 5%.

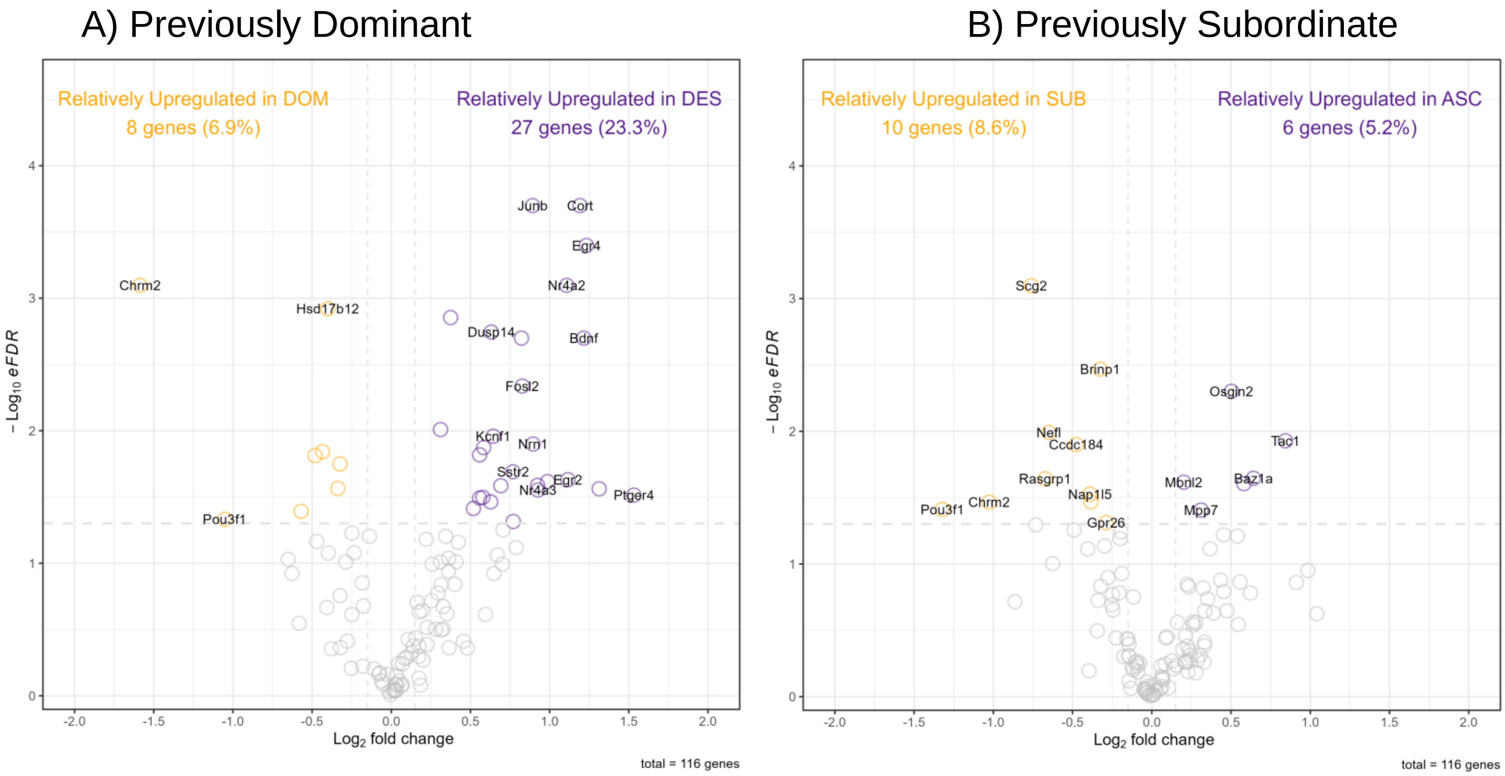

**Supplemental Figure 11:** Log normalized counts of genes related to thyroid hormone signaling in dominant animals

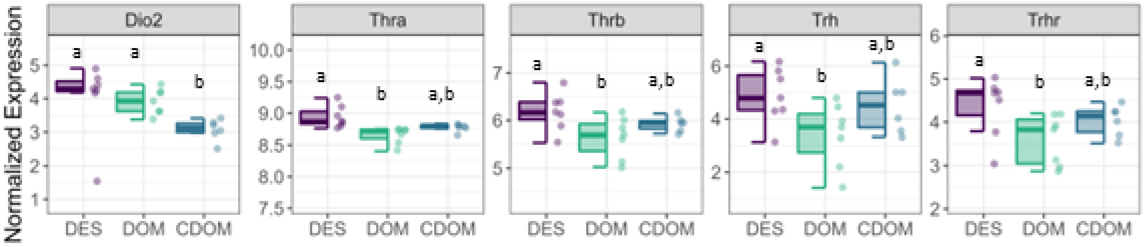

**Supplemental Figure 12:** A) Number of shared DEGs in social ascenders and descenders compared to both animals who maintained their social status and control animals. B) Median and IQR of log normalized counts of eight social transition genes differentially expressed in both descending and ascending animals. Points represent individual subjects. DOM = previously dominant males that remain dominant; DES = previously dominant males that socially descend; CDOM = control dominant animals that remain dominant. SUB = previously subordinate males that remain subordinate; ASC = previously subordinate males that socially ascend; CSUB = control subordinate animals that remain subordinate.

A)

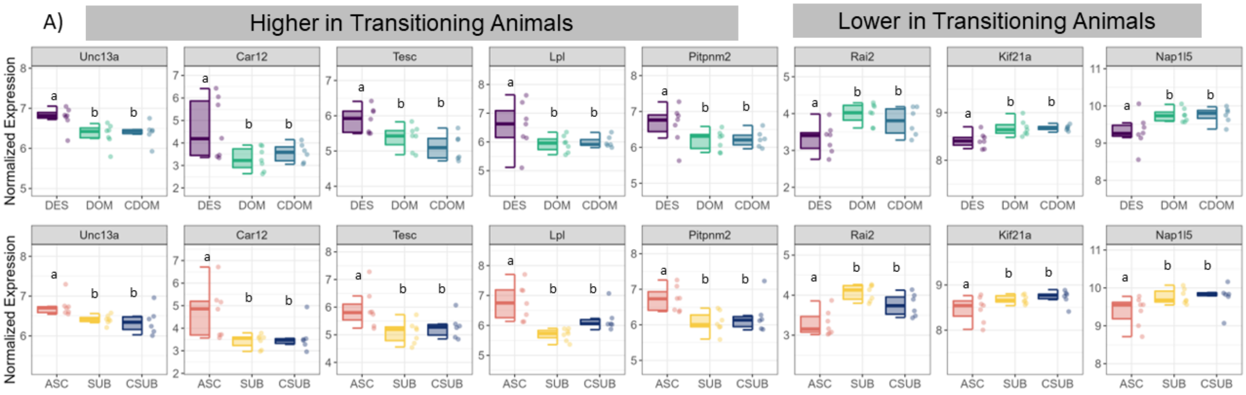


B)

| | ↑↑ ASC vs. SUB + CSUB | ↓↓ASC vs. SUB + CSUB |
|---|---|---|
| ↑↑ DES vs. DOM + CDOM | 5 | 0 |
| ↓↓ DES vs. DOM + CDOM | 0 | 3 |

**Supplemental Figure 13:** WGCNA Soft threshold plots (I) Dendrograms (II) Module correlation heatmaps (III) for all social conditions (A) and reorganized conditions (B).

A)

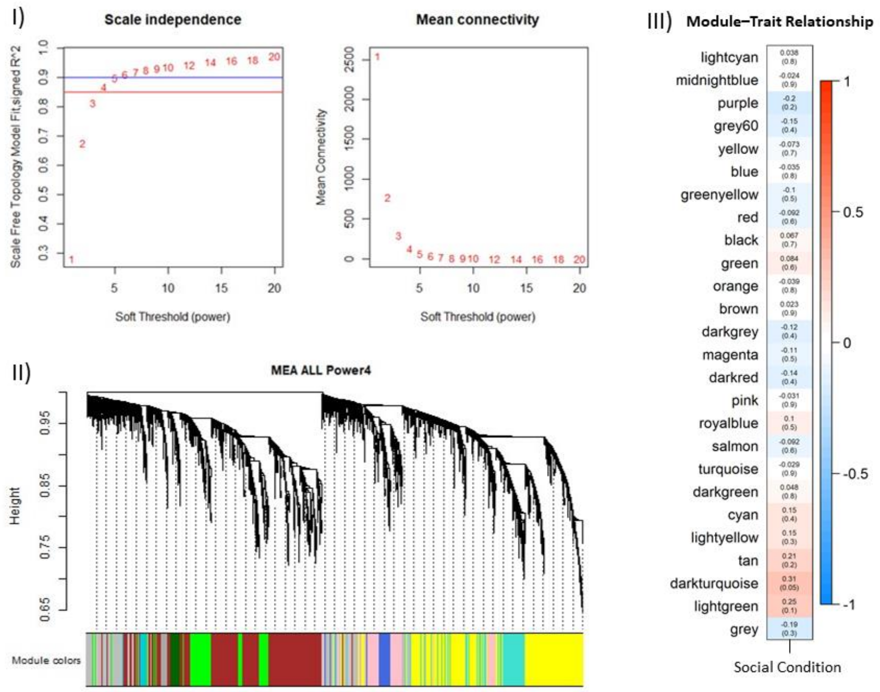


B)

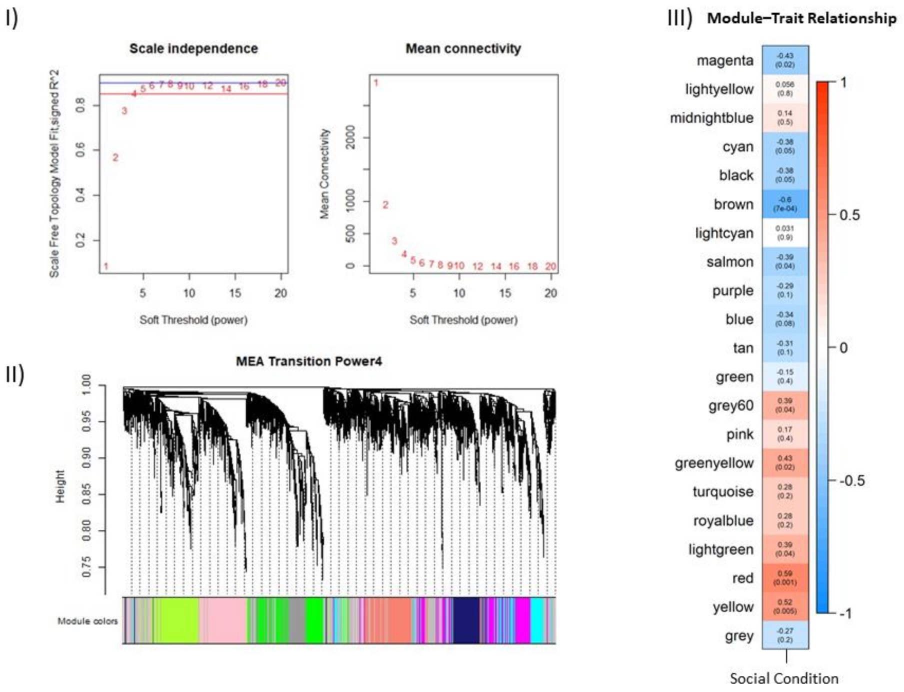

**Supplemental Table 1 -** Social Transition genes. Each gene is significantly differentially expressed in both DES vs DOM (descending animals vs animals who maintained dominant status) and ASC vs SUB (ascending animals animals vs animals who maintained subordinate status).

| symbol | DES_logFC | DES_pvalue | ASC_logFC | ASC_pvalue |
|---|---|---|---|---|
| A830018L16Rik | 0.576914 | 4.00E-04 | 0.329162 | 0.0464 |
| Abcf1 | -0.24094 | 0.0014 | -0.23414 | 0.0024 |
| Ache | -0.50638 | 0.0146 | -0.5216 | 0.0408 |
| Actn1 | 0.512624 | 0.001 | 0.463429 | 0.019 |
| Acvr1c | 1.216223 | 0.005 | 0.776506 | 0.021 |
| Adamts15 | 0.493878 | 0.0284 | 0.359582 | 0.0328 |
| Adcy2 | -0.67237 | 0.0128 | -0.57349 | 0.0342 |
| Adnp2 | 0.364177 | 0.0226 | 0.295345 | 0.0026 |
| Ak6 | -0.22997 | 0.0426 | -0.25215 | 0.0282 |
| Akap12 | -0.54412 | 0.0152 | -0.64061 | 0.0212 |
| Akr1b10 | -0.70806 | 0.0112 | -0.49523 | 0.0192 |
| Alkbh1 | 0.429255 | 0.0336 | 0.269539 | 0.019 |
| Aloxe3 | 0.801568 | 0.0066 | 0.698718 | 0.0094 |
| Ank1 | -0.65354 | 0.006 | -0.54263 | 0.0216 |
| Ano3 | 0.657585 | 0.0024 | 0.589725 | 0.0412 |
| Aplp2 | -0.25217 | 0.0022 | -0.26624 | 0.0488 |
| Arhgap33 | 0.41112 | 0.0068 | 0.645786 | 0.0076 |
| Arnt2 | -0.29393 | 0.019 | -0.22513 | 0.0402 |
| Atf7 | 0.241682 | 0.0344 | 0.252455 | 0.0208 |
| Atp1b1 | -0.21439 | 0.0166 | -0.28514 | 0.0316 |
| Atp2c1 | 0.3681 | 0.0062 | 0.303319 | 0.0022 |
| Atp6v1c2 | 0.90329 | 0.0032 | 1.42903 | 0.0012 |
| Auts2 | 0.595016 | 0.0016 | 0.478779 | 0.0134 |
| Bcor | -0.41408 | 0.0114 | -0.34416 | 0.0082 |

| | | | | |
|---|---|---|---|---|
| Bean1 | -0.35088 | 0.0122 | -0.38788 | 0.0048 |
| Bmper | -0.73108 | 0.0484 | -0.87735 | 0.007 |
| C2cd2l | 0.51791 | 4.00E-04 | 0.400995 | 0.039 |
| Car12 | 1.59098 | 0.017 | 1.217787 | 0.005 |
| Carf | 0.286358 | 0.0218 | 0.292225 | 0.011 |
| Cbln2 | -1.29416 | 0.0306 | -2.28456 | 2.00E-04 |
| Ccdc116 | 0.535357 | 0.0432 | 0.491627 | 0.029 |
| Ccdc149 | -0.24663 | 0.0422 | -0.3631 | 0.0038 |
| Ccdc163 | -0.47023 | 0.0256 | -0.48242 | 0.0422 |
| Ccdc186 | -0.31574 | 0.0092 | -0.35766 | 0.001 |
| Ccdc28b | -0.42506 | 0.0066 | -0.31516 | 0.038 |
| Cdh13 | 0.347072 | 0.0044 | 0.390885 | 0.0044 |
| Cep290 | -0.2941 | 0.014 | -0.28524 | 0.012 |
| Cep43 | -0.40281 | 4.00E-04 | -0.41888 | 0.0134 |
| Cert1 | -0.46004 | 2.00E-04 | -0.30273 | 0.0206 |
| Cgrrf1 | -0.40165 | 0.004 | -0.29726 | 0.0128 |
| Chga | -0.33121 | 0.0452 | -0.46375 | 0.0234 |
| Chrm2 | -1.58609 | 8.00E-04 | -1.02681 | 0.0342 |
| Chst11 | 0.334468 | 0.0332 | 0.49657 | 0.01 |
| Cinp | 0.26354 | 0.0388 | 0.277402 | 0.0348 |
| Cisd3 | -0.38816 | 0.0118 | -0.38069 | 0.0212 |
| Coasy | -0.33388 | 0.0038 | -0.36104 | 0.0204 |
| Col11a1 | -1.06369 | 0.009 | -0.74663 | 0.036 |
| Col23a1 | 0.953247 | 2.00E-04 | 0.82572 | 0.0296 |
| Col6a1 | 0.617155 | 0.0244 | 1.232929 | 0.0188 |
| Cpd | 0.434778 | 0.0102 | 0.353497 | 0.0278 |
| Cplane2 | 0.463229 | 0.037 | 0.484374 | 0.0342 |

| | | | | |
|---|---|---|---|---|
| Crym | 1.814335 | 0.004 | 2.09364 | 0.0068 |
| Ctps2 | -0.25534 | 0.0088 | -0.27367 | 0.0036 |
| Cxadr | 0.446209 | 0.0284 | 0.27315 | 0.0254 |
| Cybrd1 | 0.546543 | 0.0458 | 0.580657 | 0.0264 |
| Cyp26b1 | -1.43612 | 0.0274 | -1.65781 | 0.0324 |
| Dapk1 | 0.553517 | 0.005 | 0.483142 | 4.00E-04 |
| Dclk3 | 0.673757 | 0.0414 | 0.653459 | 0.0386 |
| Ddn | 1.178228 | 0.002 | 1.105207 | 0.0048 |
| Ddx1 | -0.24543 | 0.0172 | -0.2106 | 0.0262 |
| Dennd1a | 0.319858 | 0.0148 | 0.254854 | 0.004 |
| Dennd1b | 0.454137 | 0.0108 | 0.353434 | 0.0324 |
| Doc2b | 0.651758 | 0.0092 | 0.728428 | 0.0134 |
| DPCD | -0.24077 | 0.018 | -0.27905 | 0.0096 |
| Dpysl3 | -0.35217 | 0.008 | -0.43594 | 0.0288 |
| Efcab2 | -0.40986 | 0.0022 | -0.4175 | 0.0144 |
| Elavl2 | -0.4304 | 0.0228 | -0.64386 | 0.0134 |
| Endod1 | -0.55953 | 0.0016 | -0.35774 | 0.0332 |
| Erc2 | 0.368751 | 0.0444 | 0.45501 | 0.014 |
| Esrrb | -1.37009 | 0.0374 | -1.38596 | 0.044 |
| Fam124a | 0.644842 | 0.0024 | 0.500069 | 0.0198 |
| Fam126a | 0.362332 | 0.0126 | 0.529068 | 0.0034 |
| Fam184b | 0.558935 | 0.004 | 0.583888 | 0.0206 |
| Fam187b | 0.591803 | 0.0238 | 0.521903 | 0.0166 |
| Fam227a | 0.361337 | 0.04 | 0.389217 | 0.0164 |
| Fat4 | 0.978463 | 0.0286 | 0.881973 | 0.0154 |
| Fcf1 | -0.42539 | 0.0274 | -0.32471 | 0.0392 |
| Fhod3 | -0.54206 | 0.0396 | -0.67316 | 0.0038 |

| | | | | |
|---|---|---|---|---|
| Fibin | 0.919316 | 0.0048 | 0.901467 | 0.0346 |
| Fignl2 | -0.95528 | 2.00E-04 | -0.74508 | 0.048 |
| Flrt2 | -0.46333 | 0.0104 | -0.28006 | 0.035 |
| Flywch1 | -0.32962 | 0.0034 | -0.28725 | 0.0268 |
| Fundc2 | -0.44095 | 0.0018 | -0.5013 | 0.0014 |
| Fuom | -0.31081 | 0.0388 | -0.29351 | 0.037 |
| Fxyd5 | 0.460113 | 0.009 | 0.531923 | 0.0306 |
| Gda | 0.714907 | 0.0044 | 0.696404 | 0.0386 |
| Glrx3 | -0.25967 | 0.0226 | -0.24639 | 0.031 |
| Gm5176 | 0.803507 | 0.019 | 0.417904 | 0.0218 |
| Gnas | -0.20436 | 0.0092 | -0.34424 | 0.004 |
| Gpc4 | 0.787776 | 0.0384 | 0.887491 | 0.0132 |
| Gpr153 | -0.66294 | 0.0118 | -0.50162 | 0.0492 |
| Gria3 | 0.351539 | 0.0342 | 0.42329 | 0.0256 |
| Grid1 | 0.443613 | 0 | 0.254416 | 0.0046 |
| Grin2b | 0.522996 | 0.0134 | 0.428451 | 0.0384 |
| Gtpbp2 | 0.328593 | 0.0282 | 0.381475 | 8.00E-04 |
| H2-DMa | 0.658192 | 0.0016 | 0.596477 | 0.0254 |
| Haus3 | -0.58708 | 0.0024 | -0.4221 | 0.0024 |
| Heatr3 | 0.284168 | 0.0426 | 0.270441 | 0.029 |
| Hipk2 | -0.48698 | 0.0022 | -0.25815 | 0.0224 |
| Hivep1 | -0.60082 | 0.0038 | -0.33175 | 0.01 |
| Hrk | 1.067184 | 0.0028 | 0.913428 | 0.029 |
| Hsp90aa1 | -0.2872 | 0.024 | -0.32219 | 0.0012 |
| Hspa4l | -0.31339 | 8.00E-04 | -0.34043 | 0.0274 |
| Htr4 | 1.272175 | 0.004 | 1.261285 | 4.00E-04 |
| Ifitm10 | -0.597 | 0.0112 | -0.56761 | 0.0292 |

| | | | | |
|---|---|---|---|---|
| Iglon5 | -0.76236 | 0.0158 | -0.47027 | 0.021 |
| Ikzf4 | 0.446295 | 6.00E-04 | 0.326641 | 0.0456 |
| Il17rd | 0.583291 | 0.0108 | 0.497763 | 0.032 |
| Ints12 | -0.44733 | 0.043 | -0.44184 | 0.0054 |
| Itpk1 | -0.5648 | 6.00E-04 | -0.58345 | 0.0018 |
| Itpka | 1.058407 | 0.0078 | 0.970381 | 0.033 |
| Jph4 | 0.362694 | 0.0038 | 0.363882 | 0.0304 |
| Kcng1 | 0.611461 | 0.0394 | 0.487877 | 0.0388 |
| Kcnh4 | 1.359431 | 0.0206 | 1.334375 | 0.048 |
| Kcnj4 | 0.727073 | 0.0176 | 0.608933 | 0.035 |
| Kcns1 | 1.444563 | 0.0018 | 0.860838 | 0.0162 |
| Kif21a | -0.26174 | 0.0024 | -0.21934 | 0.0452 |
| Kin | -0.28857 | 0.0386 | -0.33757 | 0.0262 |
| Klhl1 | -0.98688 | 0.0126 | -1.02625 | 0.0406 |
| Klhl4 | -0.7249 | 0.0224 | -0.69851 | 0.048 |
| Lamb1 | 0.777684 | 0.0084 | 0.528155 | 0.021 |
| Laptm4b | -0.36561 | 0.011 | -0.65439 | 0.007 |
| Lcorl | 0.624974 | 0.0174 | 0.539802 | 0.0372 |
| Lhfpl3 | -0.75376 | 0.0044 | -0.40404 | 0.026 |
| Limk1 | -0.52605 | 0.0158 | -0.52866 | 0.0234 |
| Lin37 | -0.23619 | 0.0402 | -0.28398 | 0.0192 |
| Lingo3 | 0.375145 | 0.032 | 0.626091 | 0.01 |
| Lmo7 | 0.677537 | 0.0174 | 0.805429 | 0.005 |
| Lpl | 0.673022 | 0.0234 | 1.065266 | 2.00E-04 |
| Lrrc4c | 0.23567 | 0.0088 | 0.203906 | 0.0404 |
| Luzp1 | -0.5871 | 0.0012 | -0.35687 | 0.0466 |
| Magoh | -0.33315 | 0.0362 | -0.20552 | 0.0324 |

| | | | | |
|---|---|---|---|---|
| Map7d2 | -0.25013 | 0.026 | -0.31431 | 0.0496 |
| Mast3 | 0.530595 | 0.0012 | 0.446671 | 0.0126 |
| Mchr1 | 0.950051 | 6.00E-04 | 0.500846 | 0.0226 |
| Mr1 | 0.628206 | 0.0434 | 0.67835 | 0.0066 |
| Mrps18a | -0.26762 | 0.038 | -0.38989 | 0.0112 |
| Mrtfa | 0.292921 | 0.0332 | 0.344639 | 0.0254 |
| Mtmr12 | 0.653097 | 0.0138 | 0.503465 | 0.039 |
| Mxd3 | 1.481602 | 8.00E-04 | 0.95914 | 0.0034 |
| Mybpc1 | 2.952795 | 0.0124 | 2.324035 | 0.0392 |
| Mzf1 | 0.381506 | 0.0266 | 0.593283 | 0.0026 |
| Naa40 | 0.359234 | 0.0114 | 0.280414 | 0.0082 |
| Nap1l3 | -0.4735 | 2.00E-04 | -0.38543 | 0.0024 |
| Nap1l5 | -0.43397 | 0.0144 | -0.39062 | 0.0298 |
| Nars | -0.22661 | 0.0402 | -0.31341 | 0.0132 |
| Nedd4l | 0.290788 | 0.0376 | 0.237881 | 0.0462 |
| Nefl | -0.32416 | 0.0178 | -0.65017 | 0.0102 |
| Nefm | -0.72906 | 0.009 | -0.98883 | 0.017 |
| Nek10 | 0.67313 | 0.0022 | 0.6255 | 0.0042 |
| Neurl1a | 0.357858 | 0.0362 | 0.362629 | 0.0384 |
| Neurl1b | 0.63852 | 0.0038 | 0.39567 | 0.0278 |
| Nhsl2 | 0.336584 | 0.0024 | 0.252126 | 0.0276 |
| Nkrf | -0.50713 | 0.0018 | -0.35424 | 0.0212 |
| Nphp4 | 0.356758 | 0.029 | 0.41922 | 0.0134 |
| Nrbp2 | 0.333131 | 0.008 | 0.277029 | 0.0132 |
| Nrip1 | 0.473659 | 0.0326 | 0.552179 | 0.001 |
| Nsmf | 0.404612 | 0.042 | 0.468741 | 0.0266 |
| Olfr1247 | 0.399742 | 0.0194 | 0.393881 | 0.0442 |

| | | | | |
|---|---|---|---|---|
| Olfr393 | 0.634442 | 0.0224 | 0.523329 | 0.0446 |
| Olfr801 | 1.136483 | 0.0012 | 0.736895 | 0.0018 |
| Olfr806 | 1.077083 | 4.00E-04 | 0.889375 | 0.008 |
| Otulinl | 0.788781 | 0.009 | 0.557488 | 0.0128 |
| P3h3 | 0.474064 | 0.0036 | 0.423678 | 0.0368 |
| Paqr9 | 0.367383 | 0.001 | 0.339252 | 0.01 |
| Pcdh10 | -0.37534 | 0.0452 | -0.56917 | 0.0054 |
| Pcdh17 | 0.614655 | 0.0162 | 0.623496 | 0.014 |
| Pced1b | 0.383474 | 0.0322 | 0.463611 | 0.0084 |
| Pde4d | -0.41217 | 0.0064 | -0.43293 | 0.0124 |
| Pde7a | 0.298642 | 0.0388 | 0.227181 | 0.0406 |
| Peli1 | 0.453161 | 0.0014 | 0.410082 | 0.0012 |
| Pfdn2 | -0.30176 | 0.0374 | -0.30249 | 0.032 |
| Pfdn4 | -0.22671 | 0.0182 | -0.3496 | 0.0032 |
| Phlda3 | -1.03098 | 0.0014 | -0.62 | 0.0042 |
| Pitpnm2 | 0.464737 | 0.0334 | 0.632129 | 0.007 |
| Plcb4 | -0.54672 | 0.0092 | -0.47668 | 0.0442 |
| Plekhg5 | 0.410661 | 0.0296 | 0.401116 | 0.0066 |
| Plekho1 | -0.38049 | 0.0292 | -0.40098 | 0.0134 |
| Plppr4 | 0.442826 | 0.0206 | 0.43545 | 0.0362 |
| Pou3f1 | -1.05217 | 0.0466 | -1.32156 | 0.0388 |
| Ppl | 0.938337 | 0.033 | 0.739253 | 0.0424 |
| Ppp3ca | 0.415196 | 6.00E-04 | 0.442584 | 0.0124 |
| Prdm10 | 0.361716 | 0.0064 | 0.256656 | 0.0462 |
| Prdx4 | -0.41645 | 0.0346 | -0.37581 | 0.0354 |
| Prickle1 | -0.368 | 0.0154 | -0.33442 | 0.0058 |
| Prkce | 0.418845 | 0.0076 | 0.335102 | 0.0356 |

| | | | | |
|---|---|---|---|---|
| Ptpn5 | 0.261262 | 0.0166 | 0.516394 | 0.004 |
| Ptpre | 0.490026 | 0.0024 | 0.41017 | 0.004 |
| Rab40b | 0.731293 | 0.011 | 0.670945 | 0.0218 |
| Rad21 | -0.29776 | 0.002 | -0.24244 | 0.005 |
| Rai2 | -0.75914 | 0.002 | -0.78846 | 2.00E-04 |
| Rasal2 | 0.353864 | 0.01 | 0.213206 | 0.023 |
| Rdx | -0.4619 | 0 | -0.30822 | 0.0138 |
| Rfx3 | 0.890071 | 0.005 | 0.624098 | 0.0346 |
| Rgs3 | -0.61217 | 0.0058 | -0.36475 | 0.0428 |
| Rhbdl3 | 0.512565 | 0.024 | 0.35822 | 0.049 |
| Rin1 | 0.785969 | 0.0258 | 0.849802 | 0.0158 |
| Rit2 | -0.33272 | 0.0114 | -0.3939 | 0.0236 |
| Rp9 | -0.33514 | 0.0076 | -0.21521 | 0.0154 |
| Rreb1 | 1.014476 | 0.0012 | 0.922614 | 0.0448 |
| Rwdd2b | 0.564129 | 0.0302 | 0.383454 | 0.0058 |
| Scrn1 | -0.31135 | 0.0088 | -0.30854 | 0.0346 |
| Sema3c | -0.84787 | 0.0018 | -0.79478 | 0.0152 |
| Sema6d | -0.78823 | 6.00E-04 | -0.53232 | 0.01 |
| Septin6 | -0.54138 | 0.012 | -0.60273 | 8.00E-04 |
| Sfxn2 | 0.220636 | 0.0294 | 0.335181 | 0.0186 |
| Shisa7 | 0.477031 | 0.0024 | 0.416816 | 0.0376 |
| Shox2 | -1.25789 | 0.0474 | -1.49443 | 0.0338 |
| Ski | 0.309764 | 0.0212 | 0.41119 | 0.0156 |
| Slc16a2 | 0.75067 | 0.0012 | 0.65579 | 0.0362 |
| Slc25a37 | 0.503164 | 0.0014 | 0.267756 | 0.0398 |
| Slc6a3 | -1.11624 | 0.0426 | -1.61349 | 0.0124 |
| Smad3 | 0.74133 | 0.0142 | 0.722663 | 0.0272 |

| | | | | |
|---|---|---|---|---|
| Smpd3 | 0.349754 | 0.0174 | 0.433185 | 0.032 |
| Smpdl3b | 1.645859 | 0.02 | 0.889549 | 0.0438 |
| Sowaha | 0.875618 | 2.00E-04 | 0.851554 | 0.0074 |
| Sox12 | 0.245088 | 0.011 | 0.253601 | 0.027 |
| Sox4 | -0.37394 | 0.0152 | -0.27275 | 0.0138 |
| Sphk2 | -0.35872 | 0.024 | -0.34146 | 0.0024 |
| Stk32b | -0.7038 | 0.0306 | -0.96931 | 0.0308 |
| Stk39 | -0.53515 | 0.0016 | -0.47268 | 0.0094 |
| Stxbp2 | 0.355786 | 0.0014 | 0.377291 | 0.0152 |
| Styx | 0.253866 | 0.0384 | 0.370963 | 0.0092 |
| Supt16 | -0.39379 | 0.0032 | -0.27883 | 2.00E-04 |
| Sv2a | -0.32726 | 0.001 | -0.3303 | 0.0378 |
| Syt1 | -0.21959 | 0.0144 | -0.35423 | 0.009 |
| Taf7 | -0.3526 | 0.0238 | -0.41964 | 0.0044 |
| Tax1bp1 | -0.2211 | 0.0094 | -0.22313 | 0.0058 |
| Tcof1 | -0.27138 | 0.0444 | -0.26691 | 0.0176 |
| Tenm2 | 0.437471 | 0.0148 | 0.482142 | 0.0028 |
| Tesc | 0.52331 | 0.0284 | 0.837322 | 0.006 |
| Th | -1.25823 | 0.0038 | -0.88291 | 0.0084 |
| Tlk2 | -0.368 | 4.00E-04 | -0.26717 | 0.0214 |
| Tmem65 | -0.71275 | 0.02 | -1.01694 | 0.0038 |
| Tmf1 | -0.22637 | 0.033 | -0.37024 | 0.0016 |
| Tonsl | -0.72035 | 0.0014 | -0.49261 | 0.0486 |
| Trabd2b | 1.341581 | 0.0172 | 1.182776 | 0.0294 |
| Traip | 0.594975 | 0.0196 | 0.864589 | 0.0204 |
| Trhr | 0.896836 | 0.0248 | 0.960687 | 0.0194 |
| Trim12a | 0.339964 | 0.041 | 0.382985 | 0.0082 |

| | | | | |
|---|---|---|---|---|
| Trpc1 | 0.308223 | 0.034 | 0.358899 | 0.003 |
| Tspan17 | -0.64869 | 0.0136 | -0.6565 | 0.0038 |
| Tspoap1 | 0.235766 | 0.0498 | 0.329868 | 0.0152 |
| Ttc12 | -0.64165 | 0.005 | -0.61505 | 0.0302 |
| Uchl5 | -0.22014 | 0.0214 | -0.29699 | 0.0094 |
| Unc13a | 0.445289 | 0.007 | 0.329002 | 0.0176 |
| Usp6nl | -0.35053 | 0.006 | -0.24788 | 0.0228 |
| Vat1l | -0.55571 | 0.0416 | -0.40414 | 0.0404 |
| Vmn1r181 | 0.594437 | 0.022 | 0.446644 | 0.0258 |
| Vmn1r213 | 0.278584 | 0.0226 | 0.375649 | 0.0354 |
| Vps36 | -0.21551 | 0.0152 | -0.28007 | 0.028 |
| Vwc2 | -0.39979 | 0.0272 | -0.58621 | 0.0274 |
| Vwc2l | -0.78563 | 0.0038 | -0.57565 | 0.0422 |
| Wnk4 | 0.777488 | 0.026 | 1.127554 | 0.0416 |
| Zbtb1 | 0.317111 | 0.0204 | 0.25438 | 0.014 |
| Zbtb38 | -0.22866 | 0.005 | -0.24757 | 0 |
| Zcchc2 | -0.30881 | 0.049 | -0.29425 | 0.0352 |
| Zdhhc23 | 0.759351 | 0.0242 | 0.693417 | 0.0056 |
| Zdhhc5 | -0.23269 | 0.0478 | -0.29157 | 0.0242 |
| Zfp101 | -0.45087 | 0.0294 | -0.66863 | 0.0314 |
| Zfp14 | -0.56339 | 0.0106 | -0.32818 | 0.0366 |
| Zfp512b | 0.224156 | 0.0364 | 0.263564 | 0.0192 |
| Zic5 | 1.593859 | 0.001 | 1.225841 | 0.0176 |
